# Magnetoelectrics Enables Large Power Delivery to mm-Sized Wireless Bioelectronics

**DOI:** 10.1101/2023.09.01.555944

**Authors:** Wonjune Kim, C. Anne Tuppen, Fatima Alrashdan, Amanda Singer, Rachel Weirnick, Jacob T. Robinson

## Abstract

To maximize the capabilities of minimally invasive implantable bioelectronic devices, we must deliver large amounts of power to small implants; however, as devices are made smaller, it becomes more difficult to transfer large amounts of power without a wired connection. Indeed, recent work has explored creative wireless power transfer (WPT) approaches to maximize power density (the amount of power transferred divided by receiver footprint area (length × width)). Here, we analyzed a model for WPT using magnetoelectric (ME) materials that convert an alternating magnetic field into an alternating voltage. With this model, we identify the parameters that impact WPT efficiency and optimize the power density. We find that improvements in adhesion between the laminated ME layers, clamping, and selection of material thicknesses lead to a power density of 3.1 mW/mm^2^, which is over 4 times larger than previously reported for mm-sized wireless bioelectronic implants at a depth of 1 cm or more in tissue. This improved power density allows us to deliver 31 mW and 56 mW to 10-mm^2^ and 27-mm^2^ ME receivers, respectively. This total power delivery is over 5 times larger than similarly sized bioelectronic devices powered by radiofrequency electromagnetic waves, inductive coupling, ultrasound, light, capacitive coupling, or previously reported magnetoelectrics. This increased power density opens the door to more power-intensive bioelectronic applications that have previously been inaccessible using mm-sized battery-free devices.

## Introduction

More effective power delivery will allow implanted bioelectronics to support more power-intensive functions, multichannel or multimodal operations, and smaller devices. For example, recent work has shown multichannel neural recording devices that require up to 2 mW per recording channel^1–7^ and neural stimulation devices that require up to 21 mW^1,8–10^. Increasing the received power can facilitate the addition of more recording or stimulation channels. Similarly, bioelectronic devices have been developed to perform multiple functions simultaneously, enabling the monitoring and control of biological processes in multimodal closed-loop systems^7,11–14^. Each additional sensing and recording capability requires additional power. Finally, with improved wireless power transfer (WPT), implanted bioelectronics could support their existing functions in a smaller form factor that would allow for less invasive surgical implantation and access to difficult-to-reach targets^15–18^.

Many innovative approaches to WPT have been developed, yet all face trade-offs regarding depth of penetration, power density, and alignment tolerance. For example, tissue scattering and absorption of radiofrequency electromagnetic waves and light typically restrict these approaches to shallow implants. For near-field inductive coupling (NIC), power density falls considerably as devices are miniaturized, making NIC less suitable for small implants. NIC and ultrasound are very sensitive to alignment errors and are therefore difficult to implement in applications that require continuous power. Capacitive coupling and ultrasound require transmitter contact with the body, which is challenging for freely moving rodent applications or continuous operation of implants. Several reviews describe these trade-offs for wireless power transfer technologies^1,19–21^.

Magnetoelectric (ME) receivers are a promising solution for powering implantable bioelectronics because, compared to other WPT modalities, they have the potential to deliver higher power to smaller devices with better alignment tolerance and minimal signal attenuation through air or tissue^19^. While ME materials have been explored for compact antennas^22,23^, only recently have ME materials been used for WPT in bioelectronics, demonstrating up to 2 mW of power delivery^15,24–28^. The ME receivers most commonly used to power bioelectronics are multilayer laminates that convert magnetic energy into electrical energy through mechanical coupling between magnetostrictive and piezoelectric layers^15,24–28^. This conversion is most efficient when the frequency of the magnetic field matches an acoustic resonant frequency of the ME receiver, thereby generating the maximum voltage and power^26–29^.

Because ME performance depends on several material properties and this WPT approach is relatively new in bioelectronics, we hypothesized that we could increase the power density of ME receivers by optimizing their material properties. Previous research has aimed to improve ME performance by optimizing ME material properties, but this research was primarily concerned with increasing voltage generated by cm-scale ME receivers in sub-mT-scale magnetic fields^30–33^, making these findings less applicable to WPT in implantable bioelectronics. To address this limitation, we investigated how ME material properties can be optimized to improve the power transfer to mm-scale ME receivers in mT-scale magnetic fields.

Prior work has carefully analyzed optimal power delivery to ME materials under a volume constraint and we encourage readers who seek to optimize this property to refer to that work^34^. In this work, we optimize power delivery with a footprint (length × width) constraint. In most applications, a printed circuit board is typically determined based on the components footprint. Therefore, this property is often scarce when designing the planar electronics used for implantable devices. Furthermore, the footprint of the ME receiver is orders of magnitude larger than the cross sectional area (thickness × width). Thus, we define power density as the output power delivered to the load resistance divided by the footprint of the ME receiver, as has been done in prior work^35^. To achieve higher power in a small device, this power density is the factor that one would seek to optimize. In this work, we utilize longitudinal vibration mode of ME receiver and fix aspect ratio in the optimum range. The longitudinal vibration mode is the most efficient way to achieve the highest power output under safety limits and size constraints as characterized previously^25,28^. We further assume that the magnetic field is applied parallel to the length axis of the ME receivers and keep this fixed throughout the study to drive ME receivers in longitudinal vibration mode since this is the most efficient way to excite the longitudinal vibration mode.

### Approach

To improve the performance of ME receivers, we took a multi-step approach. We first reviewed prior literature and used previously reported analytical expressions and equivalent circuit models for ME receivers to determine which factors could be optimized to increase the power density (the amount of power transferred divided by receiver footprint area (length × width)). We then experimentally validated the model using an ME receiver consisting of mechanically coupled Metglas and lead zirconate titanate (PZT). Based on these studies, we identified key factors that could be experimentally controlled and optimized to increase power density. Based on these findings, we fabricated ME receivers to test if our changes indeed would improve power density experimentally. Finally, to demonstrate that ME receivers can achieve large power densities through tissue under human safety limits, we measured received power through 1-5 cm of porcine tissue.

## Results

### Equivalent Circuit Model Study

We assembled previously reported theoretical descriptions of ME coupling^29–33,36^ and investigated ME material properties that impact the induced voltage and power and can be optimized during fabrication and clamping. We use the ME voltage coefficient (α_*ME*_), which is defined as the ratio of the change in the receiver’s open-circuit voltage (OCV) to the change in the applied magnetic field^28,30–33^, as an intermediate term to validate the model and calculate the power. To calculate the received power we use the ME power coefficient (*p*_*ME*_)

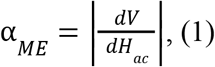

Here, *V* is the OCV from the ME receiver and *H*_*ac*_ is the applied alternating magnetic field. Note some authors prefer to define the ME voltage coefficient as the change of electric field across the piezoelectric layer induced by an applied magnetic field^23,29,37^ rather than the ME voltage coupling coefficient in Eq. 1. We choose the voltage coefficient for this work because it simplifies the calculation for power used later.

Based on the previously reported equivalent circuit model^29–33^,α_*ME*_ can be defined in terms of the magneto-elastic and electro-elastic coupling factors φ_*m*_ and φ_*p*_, respectively), the equivalent mechanical impedance (*Z*_*M*_), and the load impedance (*Z*_*L*_):

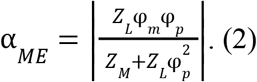

The magneto-elastic coupling factor φ_*m*_) relates the transfer of the applied magnetic field to the elastic excitation in the magnetostrictive layer (Fig. 1, green) and depends on the width (*W*), magnetostrictive layer thickness (*t*_*m*_), magneto-elastic compliance represented by tensor element *s*_33, *m*_, and piezomagnetic modulus represented by the tensor element *d*_33, *m*_ (supplementary material Eq. S2). Similarly, the electro-elastic coupling factor (φ_*p*_) relates the transfer of the mechanical stress to the electric excitation in the piezoelectric layer (Fig. 1, purple) and depends on the width (*W*), electro-elastic compliance represented by the tensor element *s*_11, *p*_, and piezoelectric modulus represented by the tensor element *d*_31,*p*_(supplementary material Eq. S3). The values of these material properties for PZT and Metglas are listed in Table S1.

**FIG. 1.**
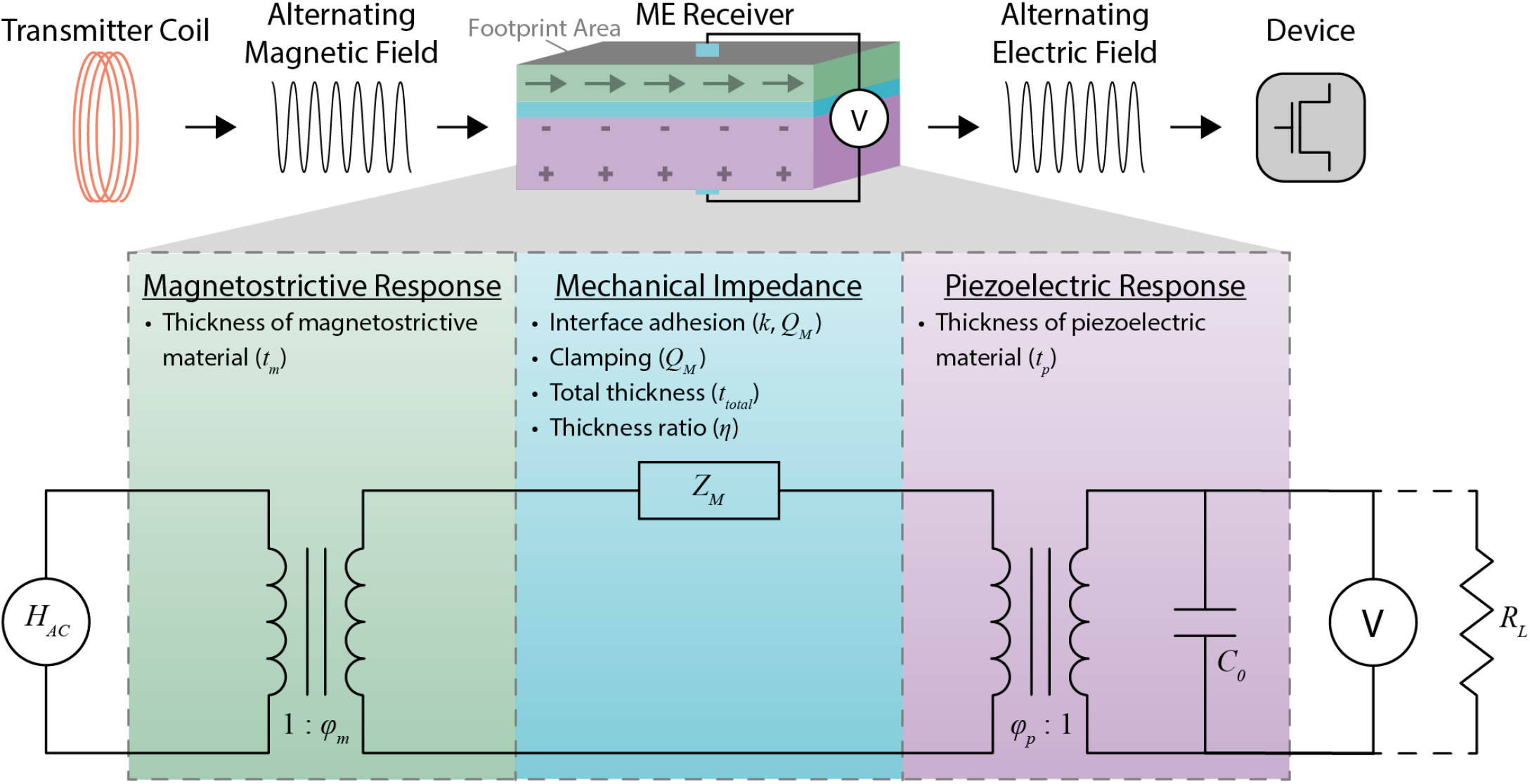
The overall mechanism and equivalent circuit model of ME WPT along with experimentally controllable ME material properties that affect the amount of power received. A transmitter coil generates an alternating magnetic field that is converted to elastic excitation in the magnetostrictive layer of the ME receiver through the magnetostrictive response (green), which is affected by the thickness of the magnetostrictive material (*t*_*m*_). Loss in the elastic excitation is expressed as mechanical impedance (blue), which is affected by the interface adhesion, clamping, total thickness (*t*_*total*_), and thickness ratio of the ME receiver (η). The interface adhesion and packaging affects both the interface coupling factor (*k*) and the mechanical quality factor (*Q*_*M*_). The clamping affects the mechanical quality factor (*Q*_*M*_). The elastic excitation is converted into an electric field through the piezoelectric response (purple), which is affected by the thickness of the piezoelectric material (*t*_*p*_). The resulting electric field can be used to wirelessly power bioelectronic devices. In the equivalent circuit model, *H*_*ac*_ is the output of the current source that represents the amplitude of the applied alternating magnetic field; φ_*m*_ and φ_*p*_ are magneto-elastic and electro-elastic coupling factors, respectively; *Z*_*M*_ is the equivalent mechanical impedance; *C*_0_ is the capacitance of the piezoelectric material; *V* is the voltage difference across the ME receiver; and *R*_*L*_ is the load resistance.

The equivalent mechanical impedance (*Z*_*M*_) depends on factors such as the interface coupling between the layers of the laminate (*k*); mechanical quality factor (*Q*_*M*_), which is a measurement of strain amplification in ME receiver; total thickness (*t*_*total*_); and thickness ratio (η), which is the ratio of the PZT thickness to total thickness (Fig. 1 and supplementary material Eqs. S4-S12). The load impedance (*Z*_*L*_) depends on the resistive load that receives the power and the capacitance of the piezoelectric layer (supplementary material Eq. S13).

To understand how α_*ME*_ can be maximized, we define the maximum ME voltage coefficient (α_*ME, max*_) as a function of *Q*_*M*_, *k, t*_*total*_, and η, which are ME material properties that can be controlled during fabrication and clamping [Table I]. To determine α_*ME, max*_, we make two assumptions. First, as depicted in Figs. 2(a-d), α depends on the driving frequency (ω = 2π*f*) of the magnetic field and is maximized when the driving frequency matches the acoustic resonant frequency (ω = ω_*r*_) of the ME receiver^28–32^. Because we are interested in peak WPT performance, we assume that the ME receivers are operating at the acoustic resonant frequency (ω_*r*_) where the voltage and received power are maximized. Additionally, we assume moderate electro-elastic coupling at the piezoelectric phase 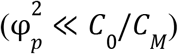 to write ω_*r*_ as a function of the average sound velocity and the length of the ME receiver (supplementary material Eq. S18). Using these assumptions and Eq. 2, we define α_*ME, max*_ as a function of *Q*_*M*_, *k, t*_*total*_, and η as follows:

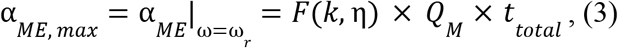

where 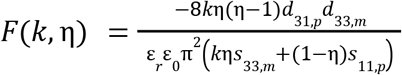 (supplementary material section 1.1). As depicted in Figs. 2(e-h), α_*ME, max*_ is linearly related to *Q*_*M*_ and *t*_*total*_ and non-linearly related to *k* and η. While α_*ME, max*_ increases with increasing *Q*_*M*_, *t* _*total*_, and k, α_*ME, max*_ is maximized at an optimal η.

**Table I.** List of the ME material properties that can be experimentally controlled and optimized.

| Name | Notation | Descriptions | Values | Controlled factors |
| --- | --- | --- | --- | --- |
| Interface coupling factor | $k$ | Mechanical coupling of Metglas and PZT interface | 0.42–0.70 | - Interface adhesion |
| Mechanical quality factor | $Q_M$ | Strain amplification in ME receiver | 43.2–75.9 | - Interface adhesion<br>- Clamping |
| Magnetostrictive material thickness | $t_m$ | Thickness of the Metglas | 25 ($\mu\text{m}$) | - Number of Metglas layers |
| Piezoelectric material thickness | $t_p$ | Thickness of the PZT | 127, 267 ($\mu\text{m}$) | - Thickness selection of PZT |
| Total thickness | $t_{total}$ | Thickness of ME receivers ($t_m + t_p$) | 152–317 ($\mu\text{m}$) | - Number of Metglas layers<br>- Thickness selection of PZT |
| Thickness ratio | $\eta$ | PZT thickness over total thickness ($t_p / t_{total}$) | 0.72–0.91 | - Number of Metglas layers<br>- Thickness selection of PZT |

**FIG. 2.**
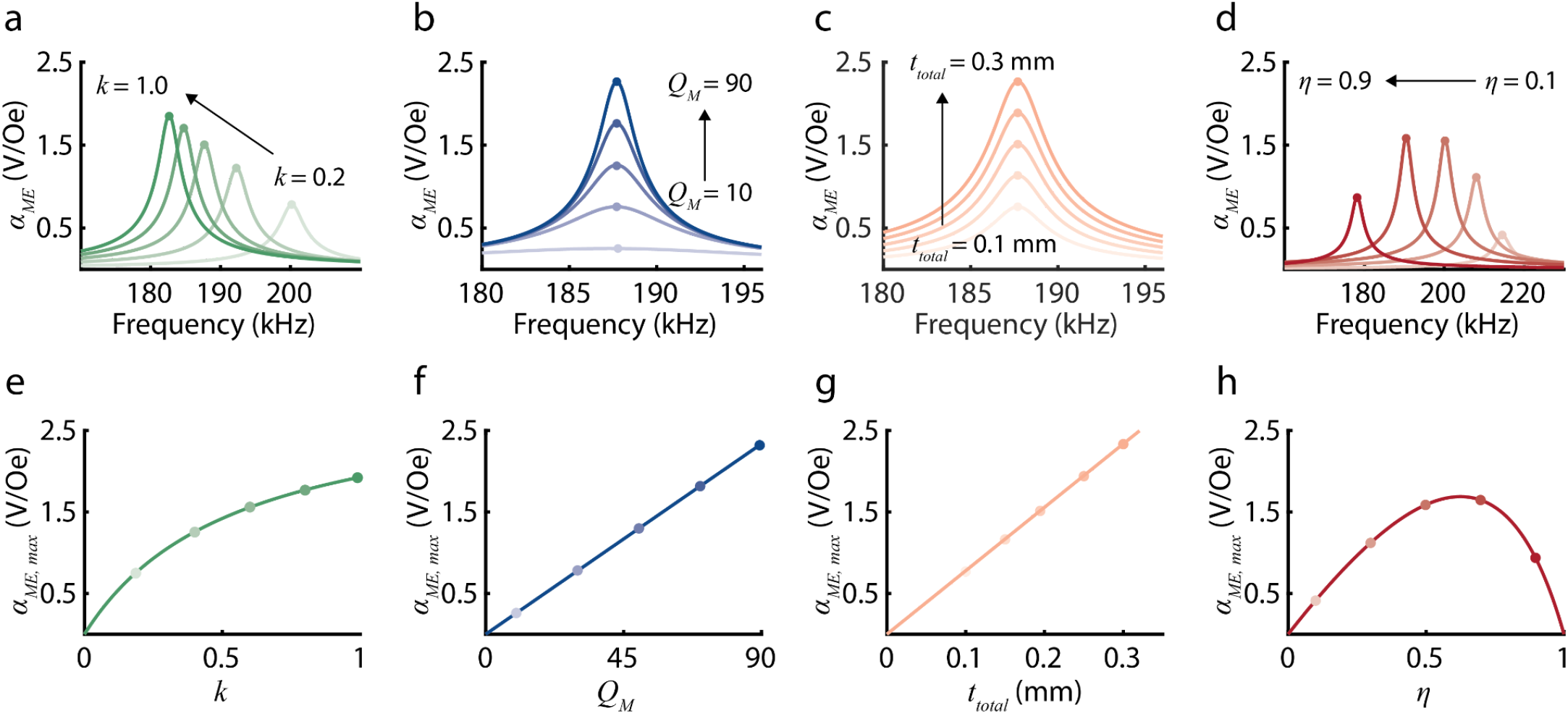
Plots show how the ME voltage coefficient (α_*ME*_) depends on several experimentally controlled variables based on our theoretical model. Panels (a-d) show α_*ME*_ as a function of frequency for varying (a) *k*, (b) *Q*_*M*_, (c) *t*_*total*_, and (d) η, which are ME material properties that can be controlled during fabrication and clamping. Increasing *k* from 0.2 to 1.0 and η from 0.1 to 0.9 decreases the resonant frequency. Increasing *Q*_*M*_ from 10 to 90 and *t*_*total*_ from 0.1 mm to 0.3 mm does not affect the resonant frequency. Panels (e-h) show the maximum ME voltage coefficient (α_*ME, max*_), which can be calculated from Eq. 3, as a function of (e) *k*, (f) *Q*_*M*_, (g) *t*_*total*_, and (h) η. α is linearly related to *Q*_*M*_ and *t*_*total*_ and non-linearly related to *k* and η. α_*ME, max*_ increases with increasing *Q*_*M*_, *t*_*total*_, and *k* and is maximized at an optimal η.

We can also quantify the ME power transfer in terms of predefined ME material properties. Here, we introduce the concept of the ME power coefficient (*p*_*ME*_). Analogous to the ME voltage coefficient, *p*_*ME*_ is defined as the ratio of the power delivered to the load (*P*_*rms*_) to the square of the amplitude of the magnetic field 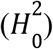:

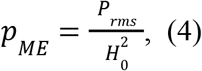

We optimize the ME power coefficient (*p*_*ME*_) to receive large power from ME receivers. Eq. 4 can be expressed in terms of other predefined variables:

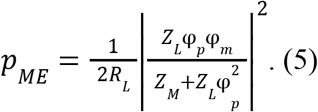

Analogous to α_*ME, max*_, we define the maximum ME power coefficient (*p*_*ME, max*_) as a function of experimentally controllable ME material properties. As depicted in Figs. 3(a) and 3(b), *p*_*ME*_ depends on load resistance (*R*_*L*_) and is maximized at an optimal load resistance 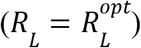 . By using the gradient descent method as reported in Ref.^29^, we can express *p*_*ME, max*_ as follows:

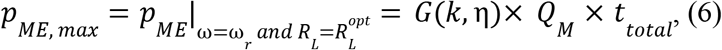

where 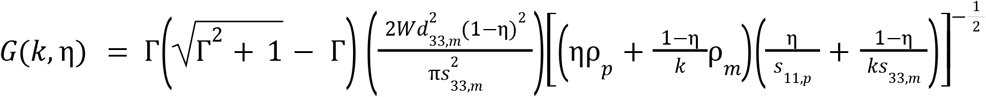 and 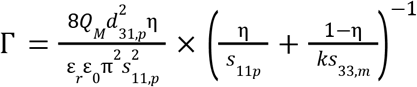 (supplementary material sections 1.2 and 1.3). This equation provides a quantitative approach to evaluating *p* in terms of these experimentally controllable properties, which we can optimize to achieve higher power density. Specifically, as depicted in Figs. 3(e-h), *p*_*ME, max*_ is linearly related to *Q*_*M*_ and *t*_*total*_ and non-linearly related to *k* and η. *p*_*ME, max*_ increases with increasing *Q*_*M*,_ *t*_*total*_, and k and is maximized at an optimal η. Authors would like to emphasize that *p*_*ME, max*_ increases linearly with *Q*_*M*_ and *t*_*total*_ . This is because the maximum output power occurs when the load matches the real (resistive) part of the equivalent ME impedance. The resistive part of the equivalent ME impedance is directly proportional to *Q* and *t* [Fig. S4(b) and (c)], and the output voltage is also linearly related to the *Q* and *t* [Fig. 3(f) and (g)]. Thus, when we calculate the output power using the expression for power 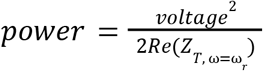, we find that power is linear with *Q* and *t* (details in supplementary material section 1.3).

**FIG. 3.**
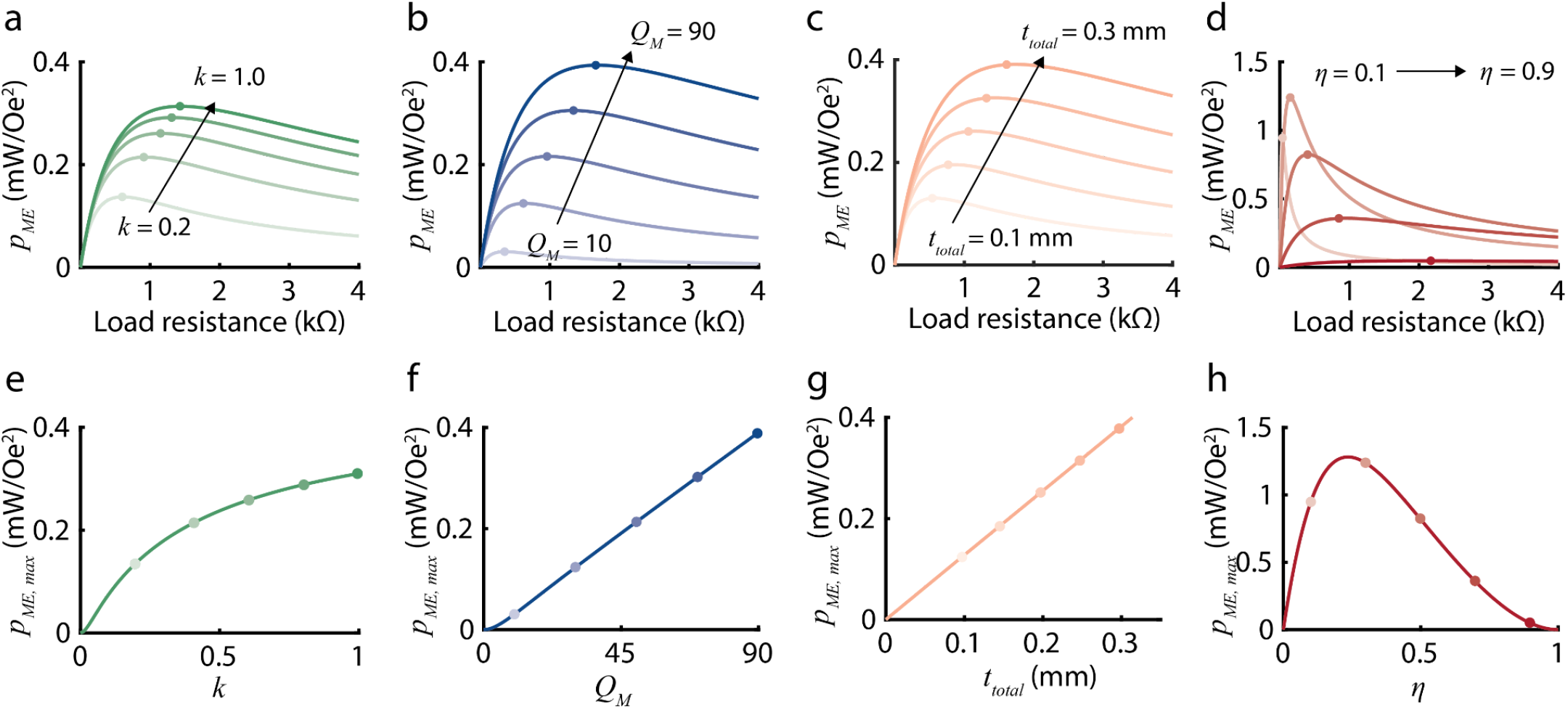
Plots show how the ME power coefficient (*p*_*ME*_) depends on several experimentally controlled variables based on our theoretical model. Panels (a-d) show *p* as a function of load resistance for varying (a) *k*, (b) *Q*_*M*_, (c) *t*_*total*_, and (d) η. Increasing *k* from 0.2 to 1.0, *Q*_*M*_ from 10 to 90, and *t*_*total*_ from 0.1 mm to 0.3 mm, and η from 0.1 to 0.9 increases the optimal load 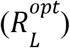 . Panel (e-h) shows the maximum ME power coefficient (*p*_*ME, max*_), which can be calculated from Eq. 6, as a function of (e) *k*, (f) *Q*, (g) *t*_*total*_, and (h) η. *p*_*ME, max*_ is linearly related to *Q*_*M*_ and *t*_*total*_ and non-linearly related to *k* and η. *p*_*ME, max*_ increases with increasing *Q*_*M*_, *t*_*total*_, and *k* and is maximized at an optimal η.

### Equivalent Circuit Model Validation

We validated the equivalent circuit model for both bilayer and trilayer configurations and for different mechanical properties of the ME receiver. Thus, we can utilize the model to accurately calculate the expected voltage and power from our ME receivers. To collect experimental data to validate our model, we fabricated 9 × 3 mm^2^ ME receivers using 267-μm-thick PZT and 25-μm-thick Metglas, measured OCV and power for each ME receiver under 0.3-mT magnetic fields, and experimentally determined *Q*_*M*_ and *k* (details in supplementary material sections 2-4). Figures 4(a) and 4(b) show theoretical and experimental values of α_*ME*_ and *p*_*ME*_ using bilayer (Metglas-PZT) and trilayer (Metglas-PZT-Metglas) ME receivers, respectively. In red, α_*ME*_ is plotted with respect to frequency, and in blue, *p*_*ME*_ is plotted with respect to load resistance. The solid lines represent theoretical values, and the circles represent experimental values. The theoretical and experimental values show close agreement for both the bilayer and trilayer configurations of the ME receiver. Figures 4(c) and 4(d) compare theoretical and experimental values of *p*_*ME, max*_ with respect to varying *Q*_*M*_ and *k*. In Fig. 4(c), red circles represent data we collected in this study using bilayer ME receivers, and a black diamond represents data from our group’s previous work^28^. As shown in Fig. 4(d), the percent error 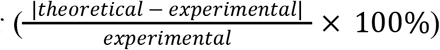 of the theoretical *p*_*ME, max*_ is less than 9% for all samples, indicating that our model is valid over a range of *Q*_*M*_ and *k*. Although we do not observe a clear functional relationship between the interface coupling coefficient (*k*) and the mechanical quality factor (*Q*_*M*_) [Fig. 4(c)], note that there may be an interdependence between these variables that may make it difficult to independently manipulate *k* and *Q*_*M*_ experimentally. Nevertheless, we analyze the effects of these two terms independently to develop an intuition as to how they separately affect the ME power transfer process.

**FIG. 4.**
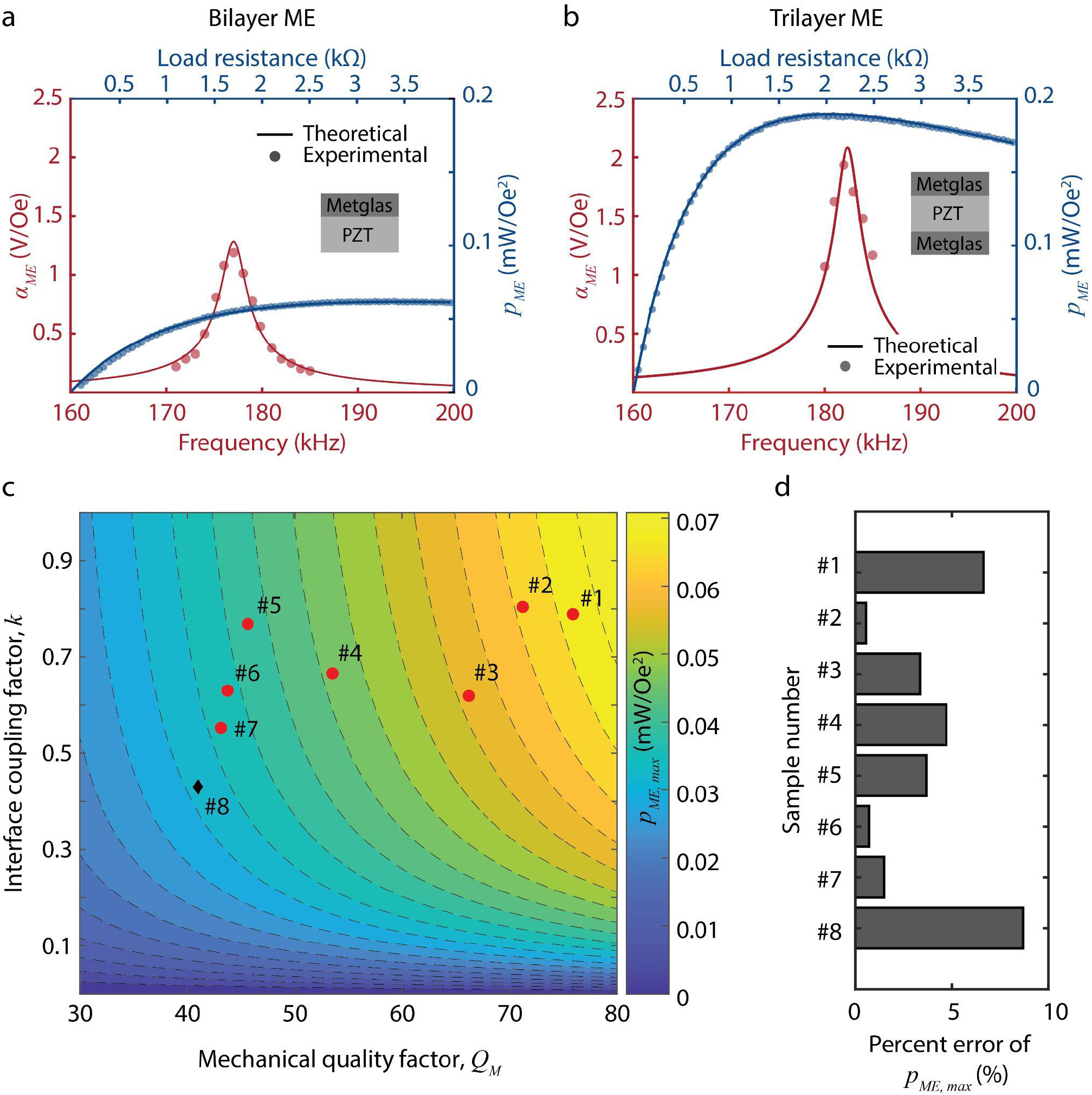
Experimental validation of the equivalent circuit model of ME WPT for both bilayer and trilayer configurations and for different mechanical properties of the ME receiver. Experimental measurements (circles) for α_*ME*_ (red) and *p*_*ME*_ (blue), show close agreement with the theoretical values (solid lines) calculated from the model for (a) bilayer (Metglas-PZT) ME receivers and (b) trilayer (Metglas-PZT-Metglas) ME receivers. (c) The 2D contour plot shows theoretical values of *p*_*ME, max*_ for varying *Q*_*M*_ and *k*. Red circles represent data we collected in this study using bilayer ME receivers, and a black diamond represents data from our group’s previous work^28^. (d) The percent error 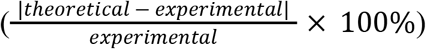 of the theoretical *p*_*ME, max*_ is less than 9% for all samples.]

### Aspect ratio optimization for power delivery under footprint area constraint

To select an aspect ratio for our ME film studies, we calculated the optimal aspect ratio (length/width, L/W) and how sensitive the power transfer was to changes in this aspect ratio. We found that at an aspect ratio of 2.5 to 3, the resulting power transfer within 97% of the peak power transfer and thus used this ratio in our experimental studies. We made this calculation by relying on the fact that the Metglas laminate exhibits a notable magnetic flux density concentration effect owing to its high permeability (μ_*r*_ = 45, 000)^37,38^.

Increasing the aspect ratio of the Metglas increases the magnetic flux density concentration, thereby resulting in an enhanced ME voltage coefficient^37^. By employing COMSOL multiphysics^38^, we simulated the magnetic flux density in a single layer of Metglas laminate as we varied the aspect ratio from 1/10 to 10/1. In this simulation, we maintained a constant footprint area and applied a magnetic field of 1 mT across an area of 27 mm2. As illustrated in Fig. 5(a) and 5(b), higher aspect ratios correlate with higher magnetic flux density within the Metglas laminate. Assuming a linear relationship between the piezomagnetic modulus (*d*_33, *m*_) of Metglas and the magnetic flux density^37^, and holding other parameters constant, we calculated normalized values for output power [Fig. 5(c)], voltage [Fig. 5(d)], and the resistive component of the ME equivalent impedance [Fig. 5(e)] relative to the aspect ratio. Notably, these calculations were conducted relative to an ME receiver with an aspect ratio of 1, as actual values would vary based on external conditions like magnetic field strength or displacement from the transmitter coil. Our analysis demonstrated an increase in output power until an optimal aspect ratio of approximately 3 (L/W=∼3) was reached. Beyond an aspect ratio of 3 the received power declined. This decline primarily stems from the increase in the resistive component of the ME impedance with the increasement of the aspect ratio [Fig. 5(e)]. Furthermore, our investigation revealed that aspect ratios within the range of 2.5 to 3 were between 97% and 99% of the optimal output power. As a result, we employ aspect ratios of 2.5 or 3 throughout this paper and mentioned the aspect ratio explicitly.

**FIG. 5.**
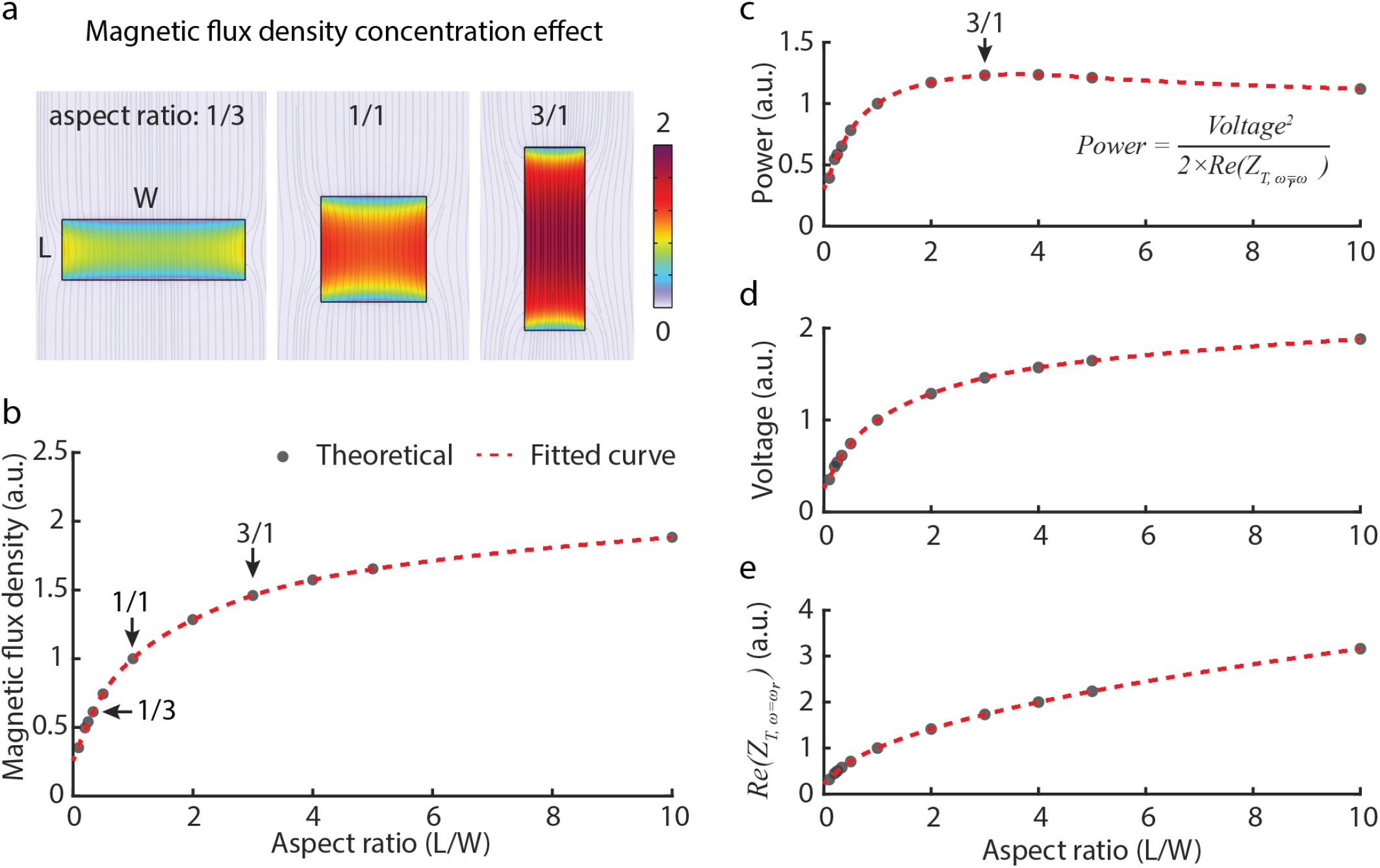
Magnetic flux density concentration effect and optimal aspect ratio for wireless power delivery within footprint constraint. (a and b) COMSOL simulation reveals magnetic flux density concentration at various aspect ratios (L/W) of the Metglas laminate. Simulated data points (black dots) span aspect ratios from 1/10 to 10/1, with a fitted curve (red dashed line) illustrating the trend. (c) Analysis highlights the maximum output power at an optimal aspect ratio of 3. This trend is primarily attributed to the (d) voltage increase resulting from magnetic flux concentration and (e) rising resistive component of the equivalent ME impedance. Aspect ratios between 2.5 and 3 are between 97% and 99% of the maximum output power. Note that simulated values in panels (b-e) are normalized to the value of aspect ratio of 1.

### Improvements in interface adhesion, clamping, and material thickness result in increased power

From our model analysis, we found three key factors that could be optimized to increase the power density of our ME receivers: interface adhesion, clamping, and selection of material thicknesses. Based on these findings, we fabricated and tested ME receivers using various methods and confirmed experimentally that these factors do indeed improve ME receiver performance as predicted.

As shown in Fig. 1 and Table I, model parameters *k, Q*_*M*_, *t*_*m*_, *t*_*p*_, *t*_*total*_, and η are ME material properties that can be optimized during fabrication and clamping to increase *p*_*ME, max*_ . Interface coupling factor (*k*) is affected by the adhesion of the Metglas and PZT layers. Mechanical quality factor (*Q*_*M*_) is affected by interface adhesion and clamping. The thicknesses of both the Metglas (*t*_*m*_) and PZT (*t*_*p*_) layers affect both total thickness (*t*_*total*_) and thickness ratio (η). Based on these dependencies, we hypothesized that we could improve power density by experimentally improving interface adhesion, clamping, and selection of material thicknesses.

We optimized interface adhesion by using a new adhesive to fabricate the ME receiver, resulting in higher output voltage. Based on previous studies, we selected seven epoxies to test: Hardman Double Bubble Red^15,25,28^, M-Bond 43-B^39^, Devcon 5 Minute^39^, West System 105A^40–42^, Epo-Tek H20E^43,44^, 3M ScotchWeld DP460^45,46^, and Masterbond EP30LV^45^. We used each epoxy to fabricate at least 90 ME receivers of 5 × 2 mm^2^ and measured the output voltage amplitude of each receiver at 1.5 mT. Figure 6(a) shows that the M-Bond epoxy results in a higher voltage amplitude compared to all other epoxies based on a Wilcoxon rank sum test (\*\*\**p*<0.001 for all pairs), while the Hardman epoxy results in the lowest voltage amplitude. The M-Bond epoxy also shows a low standard deviation in voltage amplitude compared to the other epoxies, which is a consideration for fabrication yield. Based on its high voltage amplitude and low standard deviation, we chose to use M-Bond 43-B for the subsequent studies.

**FIG. 6.**
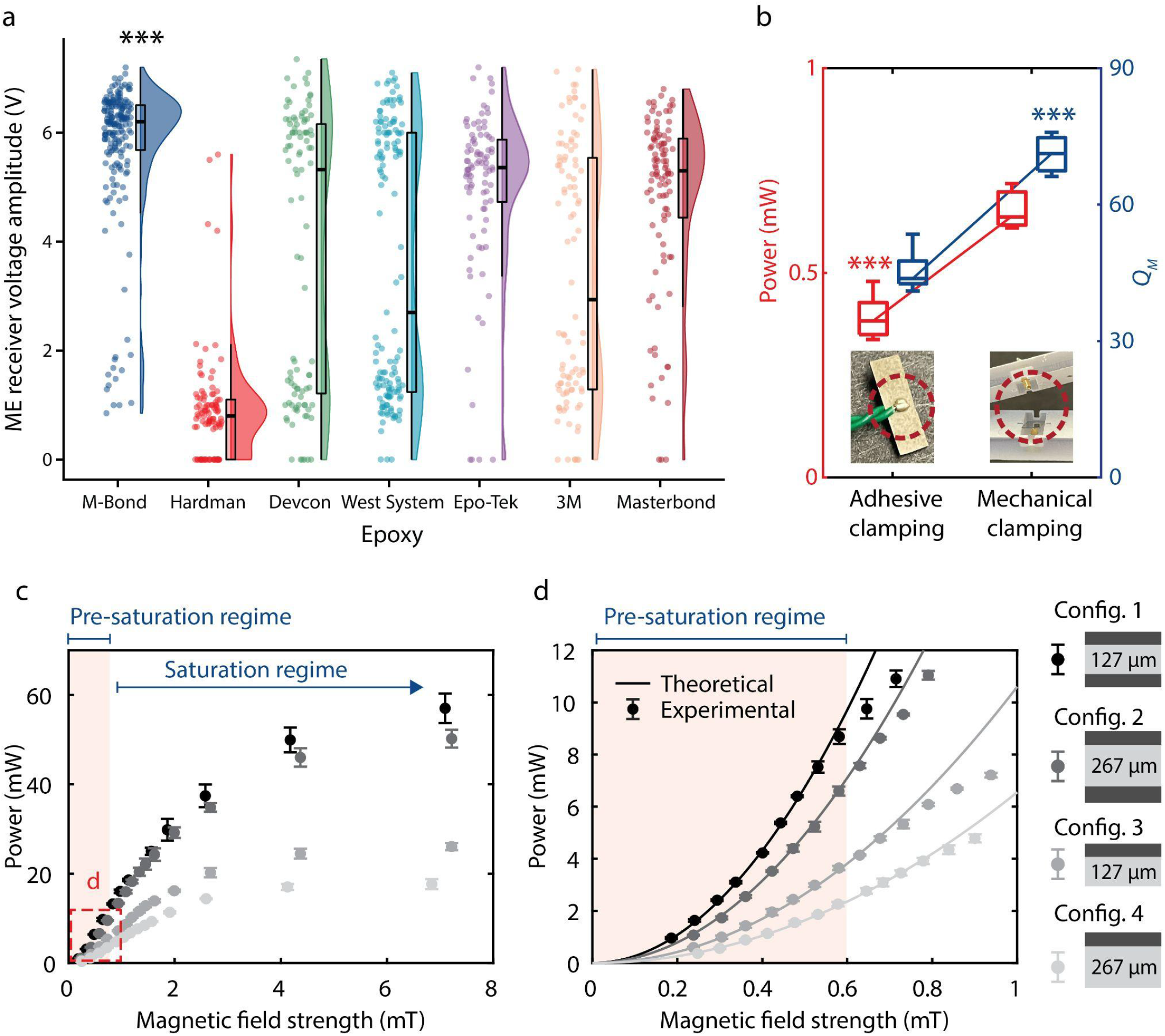
Optimizing interface adhesion, clamping, and the thickness of each layer for increased power. (a) ME receiver performance for seven different epoxies (n≥90 for each). M-Bond epoxy results in a higher voltage amplitude compared to all other epoxies (\*\*\**p*<0.001 for all pairs via Wilcoxon rank sum test). (b) Comparison of power and *Q*_*M*_ using two different clamping methods: adhesive clamping and mechanical clamping. Mechanical clamping resulted in a 67% and 57% increase in power and *Q*_*M*_, respectively (\*\*\**p*<0.001 via unpaired t-tests). (c) Measured power from four different ME receiver configurations (config. 1-4, n=3 for each) as a function of magnetic field strength. (d) Magnified version of (c) from 0-1 mT. Different shades denote different configurations. Configuration 1 exhibited the highest power, as expected from our model. The power starts saturating at 0.6 mT, resulting in lower power than theoretically predicted.

We engineered a new mechanical clamping package for the ME receiver that improved the mechanical quality factor (*Q*_*M*_) and, in turn, the received power. In prior work, an adhesive clamping method was used to connect the ME receiver to devices^15,25,28^. This method consists of applying conductive silver epoxy on the surface of the ME receiver, which results in damping of the mechanical vibrations of the receiver and accordingly a lower *Q*_*M*_ . To minimize damping and increase *Q*_*M*_, we designed a mechanical clamping method that utilizes conductive spring-loaded pins. We compared these two clamping methods using 9 × 3 mm^2^ bilayer ME receivers of 267-μm-thick PZT and 25-μm-thick Metglas under 0.3-mT magnetic fields. In Figs. 4(c) and 4(d), we used the mechanical clamping method for Samples #1 through #3, and we used the adhesive clamping method for Samples #4 through #7. Figure 6(b) shows the measured power and *Q*_*M*_ for the two clamping methods. The mechanical clamping method resulted in a 57% increase in *Q*_*M*_ and 67% increase in power (\*\*\**p*<0.001 via unpaired t-test) (details in supplementary material section 2.1).

We also increased received power by utilizing different thicknesses for the Metglas and PZT layers in our ME receivers. To test varying thickness parameters, we fabricated four versions of ME receivers (two bilayer and two trilayer configurations) (supplementary material section 3 and Table S2). We measured the maximum power received by each ME receiver (n=3) with respect to magnetic field strength [Figs. 6(c) and 6(d)]. The trilayer receiver using 127-μm-thick PZT received the largest power, 9.78 mW at 0.6 mT, which is over 250% greater than that of the bilayer ME receiver using 267-μm-thick PZT [Fig. 6(c)]. This result is consistent with the predictions we obtained from our model study. We also observed that our ME receivers begin to saturate above 0.6 mT [Fig. 6(d)]. Our model accurately predicts experimental results in the pre-saturation regime. Note that Figs. 6(c) and 6(d) compare the power at the optimal load resistance for each configuration.

### High Power Density Experimentally Demonstrated through Porcine Tissue

We demonstrated that our optimized ME receivers can achieve large power densities through tissue within human safety limits. Specifically, we transmitted power wirelessly to 5 × 2 mm^2^ and 9 × 3 mm^2^ ME receivers through *ex vivo* porcine tissue using a magnetic field of 8.0 mT at the tissue surface, which complies with the IEEE Standard C95.1-2019 electric field safety limits for humans in unrestricted environments^28^ [Fig. 7(a)]. To represent a range of implantation depths, we varied the thickness of the tissue between the transmitter and ME receiver (Tx-Rx distance) from 1 to 5 cm [Fig. 7(a) (inset)]. At each TX-RX distance, we measured the maximum power with respect to load resistance and calculated the corresponding power density. In Fig. 7(a), the red and black lines represent the 5 × 2 mm^2^ ME and the 9 × 3 mm^2^ ME receivers, respectively. The points represent the average power density of each size, and the error bars show the standard deviation. Our 5 × 2 mm^2^ and 9 × 3 mm^2^ ME receivers have power densities of 3.1 mW/mm^2^ and 2.1 mW/mm^2^, respectively, through 1 cm of tissue. With these power densities, we delivered 31 mW and 56 mW of power to the 5 × 2 mm^2^ and 9 × 3 mm^2^ ME receivers, respectively (details in supplementary material section 5).

**FIG. 7.**
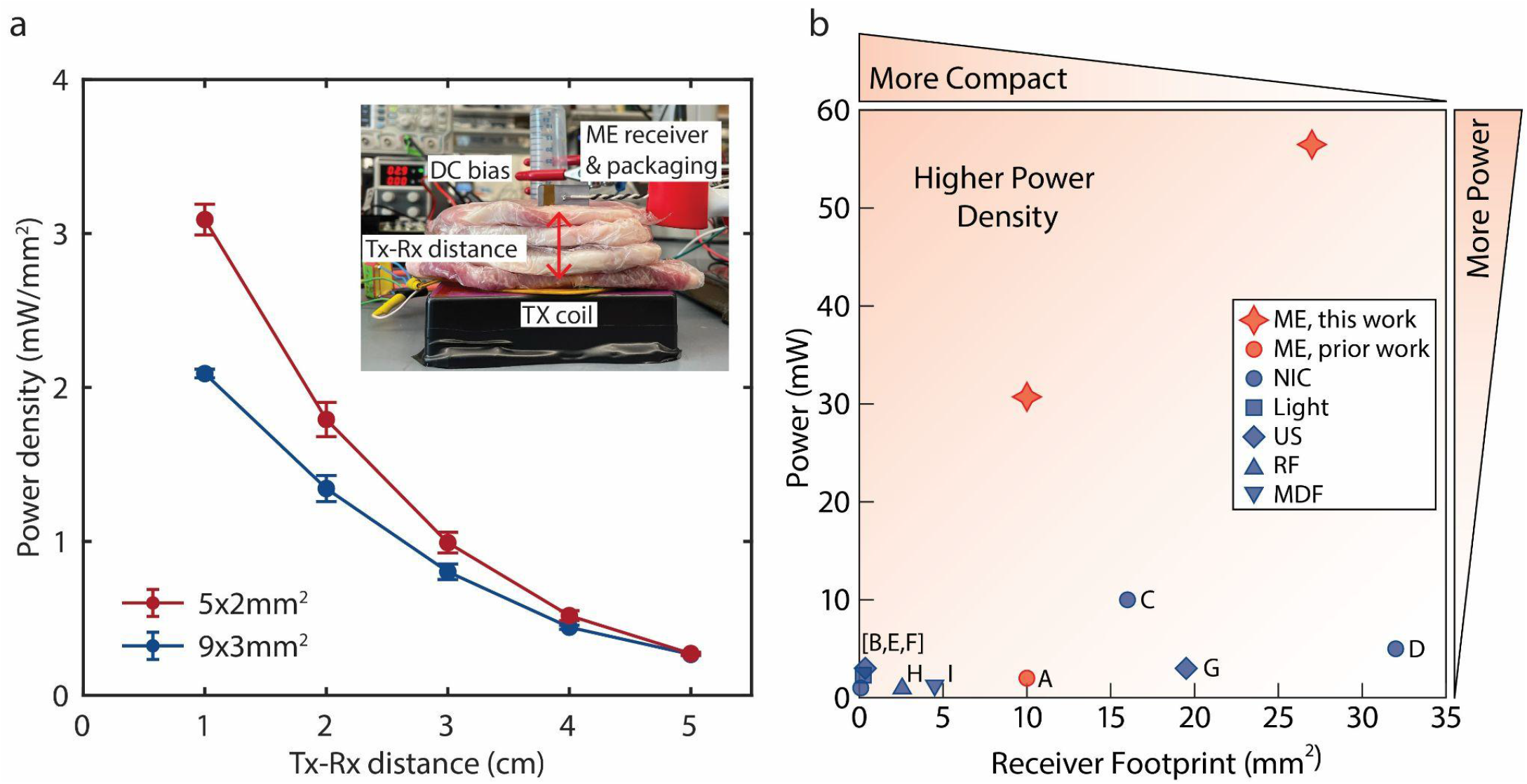
Optimized ME receivers demonstrated through *ex vivo* porcine tissue and compared to other WPT technology. (a) Measured power density in two different sizes of ME receivers as a function of distance through tissue (n=3 for each size). Through 1 cm of tissue, the power density from 5 × 2 mm^2^ and 9 × 3 mm^2^ ME receivers was 3.1 mW/mm^2^ and 2.1 mW/mm^2^, respectively, resulting in a power delivery of 31 mW and 56 mW, respectively. The inset shows the experimental setup. (b) Comparison of our ME receivers to previously reported mm-sized bioelectronics wirelessly powered by ME [A], near-field inductive coupling (NIC) [B-D], light [E], ultrasound [F, G], radiofrequency electromagnetic waves (RF) [H], and mid-field inductive coupling (MDF) [I]. Compared to a 16-mm^2^ NIC-powered device (C), we achieved over 4 times larger power density and 5 times larger power. (References : A^25^, B^47^, C^48^, D^49^, E^50^, F^51^, G^52^, H^53^, I^54^)

We compared the performance of our optimized ME receivers with similarly sized bioelectronic devices of various WPT modalities^25,47–54^ [Fig. 7(b) and Table S3]. We focused this comparison on devices compatible with clinical applications in that the transmitter power levels were within safety limits. While there are many examples of wireless power for freely moving animals, these demonstrations often use transmitter power levels that are above the IEEE safety standards for human use and/or use transmitters too large to be comfortably worn^55–58^. To directly compare the performance of different WPT modalities, we evaluated the output power and power density demonstrated in *in vivo* animal models or *ex vivo* tissue experiments with at least 1 cm Tx-Rx distance for devices with a receiver area in the range of a few tens of mm^2^. Of note, compared to a 16-mm^2^ NIC-powered device^48^, which has the largest power of previously reported devices in our comparison, we achieved over 5 times larger power.

## Discussion

In this work, we show that ME enables the largest power densities we have seen reported for mm-sized battery-free bioelectronics. We use the footprint area when calculating the power density because this metric determines the amount of space required for the power-receiving element. Thus, we believe power density based on the footprint is a relevant parameter for people who seek to integrate ME wireless power into electronic devices as described in recent publications^15,25,28^. From our equivalent circuit model analysis, we determined three key factors that could be optimized to achieve large power density: interface adhesion, clamping, and selection of material thicknesses. Our Comsol simulation analysis shows that the aspect ratio of 2.5 to 3 is within 97% of optimum power delivery. By optimizing these factors, we achieved a power of 56 mW and 31 mW through 1 cm of porcine tissue using 27-mm^2^ and 10-mm^2^ ME receivers, respectively. This is over 4 times larger power density and over 5 times larger power than previously reported for similarly sized bioelectronic devices at similar tissue depths [Fig. 7(b)]. Note that by “our” we are referring to the model we used in this manuscript which we defined based on previously reported work^29–33^.

This work can be used as a guideline for ME receiver fabrication and packaging for applications that require optimization for power, footprint area, and safety requirements. While the model we used is accurate for ME receivers only at lower magnetic field strengths (in the pre-saturation regime) [Fig. 6(d)], we find that improvements in the pre-saturation regime lead to improvements in the saturation regime even if we cannot accurately estimate those values based on our model. Additionally,the total power transferred depends onload resistance, which can be calculated from our model. As shown in Fig. 3(d), the load resistance at which the highest power is achieved depends on η. At low η, power sharply decreases at nonoptimal load resistances; at high η, power stays relatively constant at nonoptimal load resistances. These findings illustrate the importance of η and impedance matching in circuit design.

In our investigation of the optimized aspect ratio for power delivery, we operated under the assumption that the interface coupling factor and/or mechanical quality factor remained constant while varying the aspect ratio. It is important to highlight that these factors could potentially be influenced by changes in the aspect ratio, particularly due to alterations in the total length of edges. We acknowledge that a more comprehensive analysis is necessary to fully grasp the impact of aspect ratio on power delivery. The interface adhesion, clamping, and selection of material thicknesses could be further improved in future work. Of all the epoxies we tested, the M-Bond epoxy was the only one-part epoxy and had the lowest viscosity, which may be why the M-Bond epoxy resulted in receivers with the most consistently high voltage. Future work could include comparing various other one-part, low-viscosity epoxies or developing a new epoxy. Additionally, we used a brush to manually apply the M-Bond epoxy, which offers little control of the epoxy thickness and could result in inconsistent epoxy thicknesses across one receiver or between receivers. Alternative adhesive application methods to explore in future work for improved control and consistency of the epoxy layer thickness include spin-coating, spraying, or rolling. Furthermore, prior studies have shown varying results for how properties of the epoxy layer affect the ME voltage coefficient^39–43^, so future work could further examine how various epoxy layer properties affect the power of ME receivers. Another challenge to solve in future work is miniaturization and incorporation of the clamping method introduced in this work into implantable devices. The spring-loaded pins in the mechanical clamping design occupy more vertical space than the previously used conductive silver epoxy, which is a trade-off to consider when designing an implant. Additionally, packaging ME receivers into safely implantable devices will involve the selection of biocompatible materials and extended lifetime testing. Lastly, when testing material thicknesses, we used Metglas and PZT that are commercially available, which limited our design options. Future work could explore fabricating Metglas and PZT at custom thicknesses or utilizing other magnetostrictive and piezoelectric materials.

This work shows that ME WPT is heavily dependent on the receiver’s material properties; therefore, advancement and innovation in ME material engineering may further improve the power density and total power transfer of ME receivers. Here, we showed an order of magnitude improvement when compared to previously reported ME recievers, but there may be many more opportunities to increase this further with additional developments in materials design, fabrication, and packaging. Indeed the material science of ME WPT may unlock even more efficient methods to power mm-sized bioelectronics.

## Supplementary Material

See the supplementary material for detailed equations, parameter determinations, ME receiver configurations, experimental setup, *ex-vivo* experiment, and comparison with other wireless power transfer modalities.

## Acknowledgments

The authors thank Ellie Chen, Josh Chen, Abdeali Dhuliyawalla, Matthew Parker, and Josh Woods for useful discussions on magnetoelectrics and visualization. The authors thank Boshuo Wang, Zhongxi Li, Angel Peterchev, and Stefan Goetz from Duke University for providing driver electronics for the transmitter.

This research was developed with funding in part from the Robert and Janice McNair Foundation, McNair Medical Institute, the National Institutes of Health NIH grant no. U18EB029353, and the Defense Advanced Research Projects Agency (DARPA), Contract No. FA8650-21-2-7119. The views, opinions and/or findings expressed are those of the authors and should not be interpreted as representing the official views or policies of the Department of Defense or the U.S. Government.

The authors used the following tools to revise their original text: Chat GPT, Wordtune, and Grammarly. Figure 5(a) was created using the Shiny app RainCloudPlots. Figure S5 was created in part with BioRender.com.

## Author Contributions

**Wonjune Kim:** Conceptualization (supporting); Data curation (lead); Formal analysis (equal); Methodology (equal); Project administration (supporting); Software (equal); Validation (equal); Visualization (equal); Writing – original draft (supporting); Writing – review and editing (supporting). **C. Anne Tuppen:** Conceptualization (supporting); Data curation (supporting); Formal analysis (equal); Methodology (equal); Project administration (supporting); Software (equal); Validation (equal); Visualization (equal); Writing – original draft (lead); Writing – review and editing (supporting). **Fatima Alrashdan:** Formal analysis (equal); Methodology (supporting); Writing – review and editing (supporting). **Amanda Singer:** Data curation (supporting); Formal analysis (supporting); Methodology (supporting); Validation (supporting); Writing – review and editing (supporting). **Rachel Weirnick:** Data curation (supporting); Formal analysis (supporting); Software (equal); Validation (supporting); Writing – review and editing (supporting). **Jacob T. Robinson:** Conceptualization (lead); Funding acquisition (lead); Project Administration (lead); Resource (lead); Supervision (lead); Visualization (supporting); Writing – review and editing (lead).

