## Supplementary Material for "Magnetoelectrics Enables Large Power Delivery to mm-Sized Wireless Bioelectronics"

### Section 1. Model study

#### 1.1. Magnetoelectric voltage coefficient ($\alpha_{ME}$) calculation

We utilized previously reported equivalent circuit models[1–6] to derive a simplified expression for the ME voltage coefficient as a function of the parameters that we can optimize. First, as noted in Eq. 1 and Eq. 2, the ME voltage coefficient can be defined as the ratio of the change in the receiver's open circuit voltage (OCV) to the change in the applied magnetic field and expressed as follows:

$$\alpha_{ME} = \left|\frac{dV}{dH_{ac}}\right| = \left|\frac{Z_L \varphi_m \varphi_p}{Z_M + Z_L \varphi_p^2}\right|, \text{(S1)}$$

Where magneto-elastic coupling factor ($\varphi_m$) and electro-elastic coupling factor ($\varphi_p$) are a function of material properties as well as the geometry of the ME receiver. These coupling factors can be expressed as:

$$\varphi_m = W t_m \frac{d_{33,m}}{s_{33,m}}, \text{(S2)}$$

$$\varphi_p = - W \frac{d_{31,p}}{s_{11,p}}, \text{(S3)}$$

where, $d_{33,m}$ and $d_{31,p}$ are the piezomagnetic or the piezoelectric modulus, respectively, and $s_{33,m}$ and $s_{11,p}$ are the magneto-elastic or the electro-elastic compliance, respectively. Note that the values of these material properties for lead zirconate titanate (PZT) and Metglas are listed in Table SI.

[Table SI. Material properties of Metglas and PZT.]

| | Metglas | PZT |
|---|---|---|
| Material density kgm$^{-3}$ ($\rho_m$, $\rho_p$) | $7.18\times10^3$ | $7.87\times10^3$ |
| Relative permeability/permittivity ($\mu_r$, $\varepsilon_r$) | 45,000 | 1,800 |
| Piezomagnetic/piezoelectric modulus mA$^{-1}$, mV$^{-1}$ ($d_{33,m}$, $d_{31,p}$) | $9.09\times10^{-12}$ | $15.1\times10^{-12}$ |
| Electroelastic/magnetoelastic compliance m$^2$N$^{-1}$ ($s_{33,m}$, $s_{11,p}$) | $11.8\times10^{-9}$ | $-190\times10^{-12}$ |

The equivalent mechanical impedance ($Z_m$) can be expressed as follows:

$$Z_M = R_M + j\omega L_M + \frac{1}{j\omega C_M}. \text{ (S4)}$$

Under the resonant condition, each component of the $Z_m$ can be expressed as:

$$R_M = \frac{\pi Z_0}{8Q_M},\ L_M = \frac{\pi Z_0}{8\omega_0},\ C_M = \frac{8}{\pi Z_0\omega_0}, \text{ (S5)}$$

Where, characteristic frequency (i.e. closed-circuit resonant frequency, $\omega_0$) and characteristic impedance ($Z_0$) can be calculated by the formula:

$$\omega_0 = \pi\frac{\bar{v}}{L}, \text{ (S6)}$$

$$Z_0 = A\overline{v\rho}, \text{ (S7)}$$

Additionally, average density ($\bar{\rho}$), average sound velocity ($\bar{v}$), cross-sectional area ($A$), total thickness ($t_{total}$), and thickness ratio ($\eta$) are defined as follows:

$$\bar{\rho} = \eta\rho_p + \frac{(1-\eta)}{k}\rho_m, \text{ (S8)}$$

$$\bar{v}^2 = \frac{1}{\bar{\rho}}\left(\frac{\eta}{s_{11,p}} + \frac{1-\eta}{ks_{33,m}}\right), \text{ (S9)}$$

$$A = Wt_{total}, \text{ (S10)}$$

$$t_{total} = t_p + t_m, \text{ (S11)}$$

$$\eta = \frac{t_p}{t_{total}}. \text{ (S12)}$$

Note that we define the mechanical quality factor ($Q_M$) and interface coupling factor ($k$) in the same manner as discussed in Refs [1,7]. The load impedance ($Z_L$) can be represented as a parallel connection of the load resistor $R_L$ and PZT capacitance ($C_0$) as:

$$Z_L = R_L || \frac{1}{j\omega C_0} = \frac{R_L}{j\omega C_0 R_L + 1}, \text{(S13)}$$

where the capacitance of the PZT can be calculated with relative permittivity ($\varepsilon_r$), vacuum permittivity ($\varepsilon_0$), width ($W$), length ($L$), thickness of PZT($t_p$) as:

$$C_0 = \varepsilon_r \varepsilon_0 \frac{WL}{t_p}. \text{(S14)}$$

For open circuit conditions where $R_L \to +\infty$, $Z_L$ can be written as:

$$Z_L \simeq \frac{1}{j\omega C_0}. \text{(S15)}$$

By substituting $Z_L$ and $Z_m$ in the Eq. S1 with the Eq. S4 and Eq. S15 we can express $\alpha_{ME}$ with the component in the equivalent circuit model under driving frequency ($\omega$) as follows:

$$\alpha_{ME} = \left| \frac{Z_L \varphi_m \varphi_p}{Z_m + Z_L \varphi_p^2} \right| = \left| \frac{1}{j\omega C_0} \right| \left| \frac{\varphi_m \varphi_p}{R_m + j\omega L_m + \frac{1}{j\omega C_m} + \frac{\varphi_p^2}{j\omega C_0}} \right|. \text{(S16)}$$

Now, we focus on the ME voltage coefficient at the acoustic resonant frequency (i.e. open-circuit resonant frequency) of the ME receivers ($\omega_r$). As discussed in previous studies[1–5,7] and depicted in Fig. 2a-d, $\alpha_{ME}$ reaches its maximum when the driving frequency of the magnetic field matches the acoustic resonant frequency of the ME receiver ($\omega_r$). $\omega_r$ is determined by:

$$\left| j\omega_r L_M + \frac{1}{j\omega_r C_M} + \frac{\varphi_p^2}{j\omega_r C_0} \right| = 0. \text{(S17)}$$

By rearranging the above equation in terms of $\omega_r$ we obtain:

$$\omega_r = \left[ \frac{1}{L_M} \left( \frac{1}{C_M} + \frac{\varphi_p^2}{C_0} \right) \right]^{1/2}. \text{(S18)}$$

Although the value of acoustic resonant frequency ($\omega_r$) is distinct from the value of characteristic frequency ($\omega_0$), under the condition of moderate electro-elastic coupling at the piezoelectric phase$\left(\varphi_p^2 \ll C_0/C_M\right)$, we can approximate $\omega_r$ to $\omega_0$ as:

$$\omega_r \simeq \omega_0 = \pi \frac{\bar{\nu}}{L}. \text{(S19)}$$

Now, plugging in the approximation of Eq. S19 into the Eq. S16, we can obtain the maximum ME voltage coefficient at the resonant frequency as:

$$\alpha_{ME,\,max} = \alpha_{ME}\big|_{\omega=\omega_r} \simeq \alpha_{ME}\big|_{\omega=\omega_0} = \left|\frac{1}{j\omega_0 C_0}\right|\left|\frac{\varphi_m\varphi_p}{R_M + j\omega_0 L_M + \frac{1}{j\omega_0 C_M} + \frac{\varphi_p^2}{j\omega_0 C_0}}\right|. \text{ (S20)}$$

By substituting the variables in Eq. S20 with the expressions from Eq. S2-S3 and S5-S12, we obtain a simplified expression of $\alpha_{ME,\,max}$ as:

$$\alpha_{ME,\,max} = \frac{-8k\eta(\eta-1)d_{31,p}d_{33,m}Q_M t_{total}}{\varepsilon_r\varepsilon_0\pi^2\left(k\eta s_{33,m} + (1-\eta)s_{11,p}\right)}. \text{ (S21)}$$

We validated our simplified model of $\alpha_{ME,\,max}$ (Eq. S21) by comparing it with the original model. In Fig. S1, the black solid lines represent the $\alpha_{ME}$ calculated from the original model using Eq. S16 and taking the maximum values, while the red dashed lines represent the $\alpha_{ME,\,max}$ calculated from the simplified model using Eq. S21. Our comparisons show that the simplified model closely agrees with the original model in the range of $0 < k < 1$ (Fig. S1a), $0 < Q_M < 90$ (Fig. S1b), $0\,mm < t_{total} < 0.35\,mm$ (Fig. S1c), and $0 < \eta < 1$ (Fig. S1d).

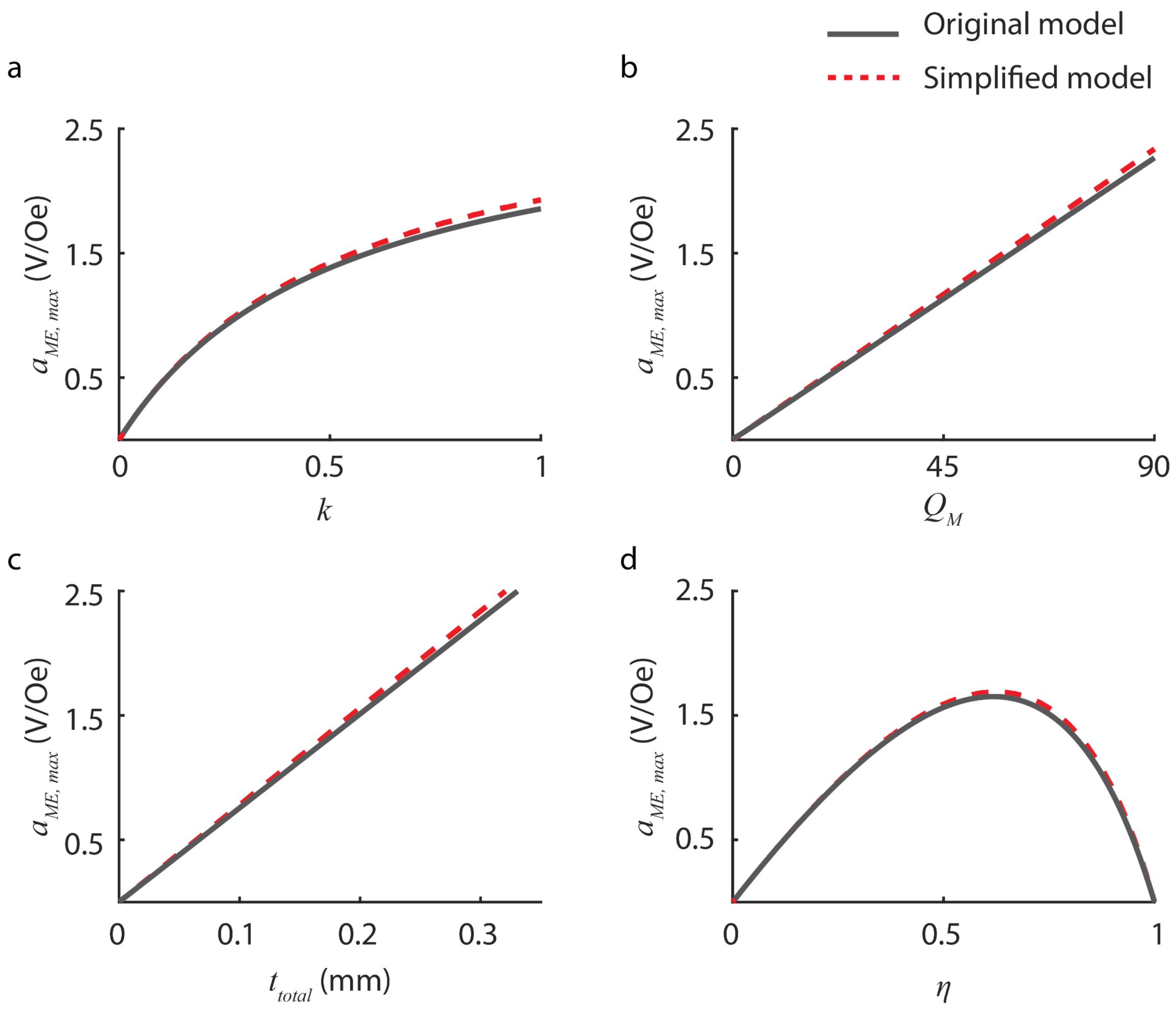


[FIG. S1. Comparison of $\alpha_{ME,\,max}$ using the original model (black solid lines) and simplified model (red dashed lines). Values calculated from the simplified model closely agree with ones calculated from the original model in the range of (a) $0 < k < 1$, (b) $0 < Q_M < 90$, (c) $0\,mm < t_{total} < 0.35\,mm$, and (d) $0 < \eta < 1$]

#### 1.2. Magnetoelectric power coefficient ($p_{ME}$) calculation

We take a similar approach as introduced in section 1.1 to calculate the ME power coefficient ($p_{ME}$) and derive a simplified expression for the $p_{ME}$ as a function of the parameters that we can optimize. The amplitude of the output voltage across the load resistor is calculated as:

$$V_0 = \left| \frac{Z_L \varphi_p \varphi_m}{Z_M + Z_L \varphi_p^2} \right| \times H_0. \text{ (S22)}$$

Note that in the power calculation, the load resistance is a finite number, so we cannot use the approximation in Eq. S15. The root means squared power delivered to the load resistance is derived as follows:

$$P_{rms} = \frac{V_0^2}{2R_L} = \frac{1}{2R_L}\left|\frac{Z_L\varphi_p\varphi_m}{Z_M+Z_L\varphi_p^2}\right|^2 \times H_0^2, \text{(S23)}$$

From the definition, $p_{ME}$ is expressed as:

$$p_{ME} = \frac{1}{2R_L}\left|\frac{Z_L\varphi_p\varphi_m}{Z_M+Z_L\varphi_p^2}\right|^2. \text{(S24)}$$

By substituting $Z_m$ and $Z_L$ in the Eq. S24 with the Eq. S4, S5, and S13 we can express $p_{ME}$ with the component in the equivalent circuit model under the driving frequency (ω) as follows:

$$p_{ME} = \frac{1}{2R_L}\left|\frac{R_L}{j\omega C_0R_L+1}\frac{\varphi_p\varphi_m}{R_M+j\omega L_M+\frac{1}{j\omega C_M}+\frac{\varphi_p^2}{j\omega C_0}+\frac{R_L\varphi_p^2}{j\omega C_0R_L+1}}\right|^2. \text{(S25)}$$

We can calculate the maximum ME power coefficient at the acoustic resonant frequency ($\omega_r$) and optimal load resistance ($R_L^{opt}$). Optimal load resistance is where the load impedance matches the output impedance of the ME receiver, which will be further discussed in the following section. By substituting variables in Eq. S25 with the expression from Eq. S2-S3 and S5-S12 and using the gradient descent method[1], we obtain $p_{ME,\,max}$ as:

$$p_{ME,\,max} = p_{ME}|_{\omega=\omega_r \text{ and } R_L=R_L^{opt}} = G(k,\eta)\times Q_M \times t_{total}, \text{(S26)}$$

where $G(k,\eta) = \Gamma\left(\sqrt{\Gamma^2+1}-\Gamma\right)\left(\frac{2Wd_{33,m}^2(1-\eta)^2}{\pi s_{33,m}^2}\right)\left[\left(\eta\rho_p+\frac{1-\eta}{k}\rho_m\right)\left(\frac{\eta}{s_{11,p}}+\frac{1-\eta}{ks_{33,m}}\right)\right]^{-\frac{1}{2}}$, and $\Gamma = \frac{8Q_Md_{31,p}^2\eta}{\varepsilon_r\varepsilon_0\pi^2s_{11,p}^2}\times\left(\frac{\eta}{s_{11p}}+\frac{1-\eta}{ks_{33,m}}\right)^{-1}$. Here, $G(k,\eta)$ is a function of $k$ and η but independent of the $t_{total}$ and can be approximated as constant with respect to $Q_M$.

We validated our simplified model of $p_{ME,\,max}$ (Eq. S26) by comparing it with the original model in the same manner we did in section 1.1. In Fig. S2, the black solid lines represent the $p_{ME}$ calculated from the original model using Eq. S25 and taking the maximum values, while the red dashed lines represent the $p_{ME,\,max}$ calculated from the simplified model using Eq. S21. Our comparisons show that the values from the simplified model closely agree with ones from the original model in the range of $0 < k < 1$ (Fig. S2a), $0 < Q_M < 90$ (Fig. S2b), $0\,mm < t_{total} < 0.35\,mm$ (Fig. S2c), and $0 < \eta < 1$ (Fig. S2d).

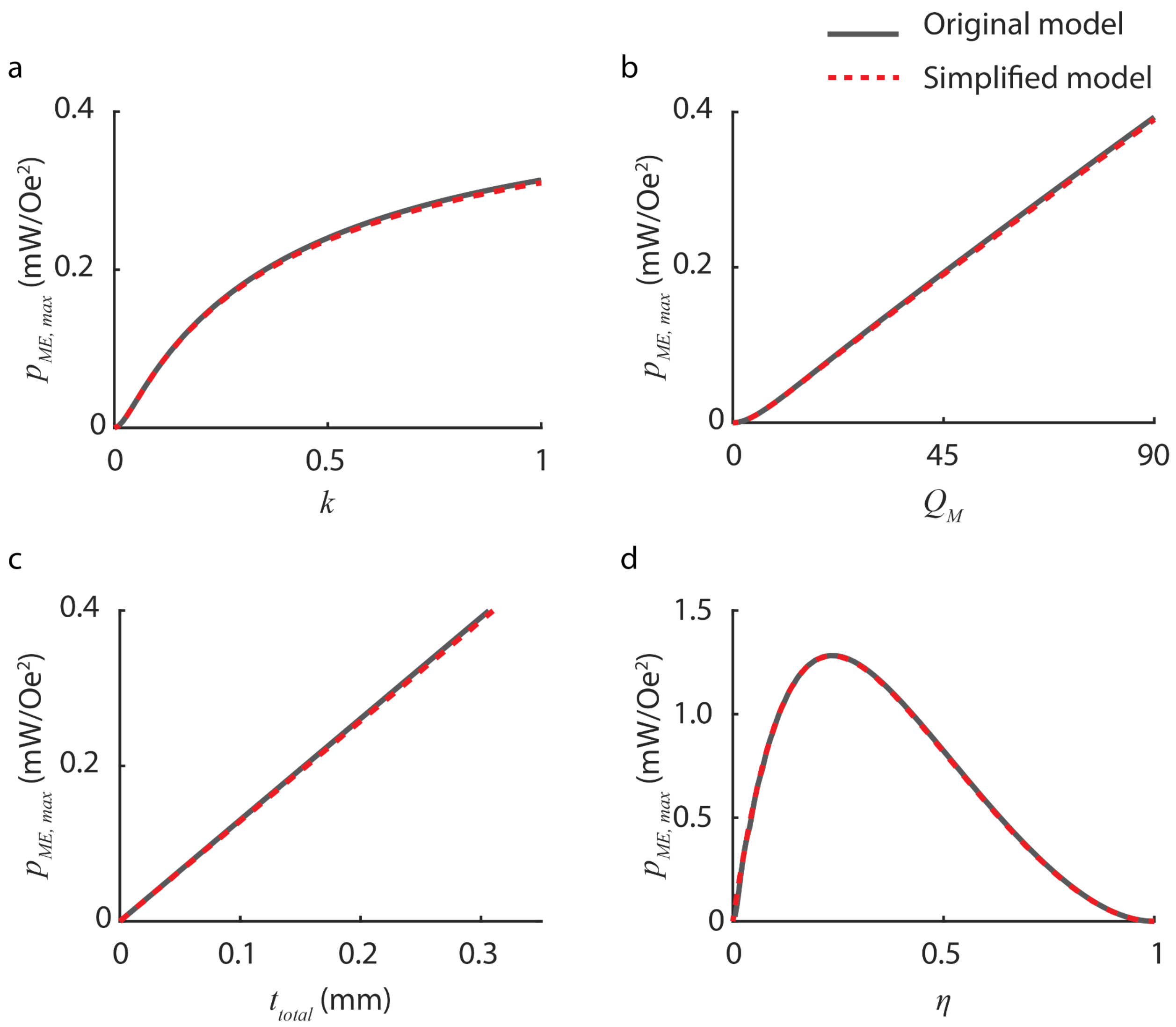


[FIG. S2. Comparison of $p_{ME, max}$ using the original model (black solid lines) and simplified model (red dashed lines). Values calculated from the simplified model closely agree with ones calculated from the original model in the range of (a) $0 < k < 1$, (b) $0 < Q_M < 90$, (c) $0\, mm < t_{total} < 0.35\, mm$, and (d) $0 < \eta < 1$]

**1.3. Optimal load resistance calculation**

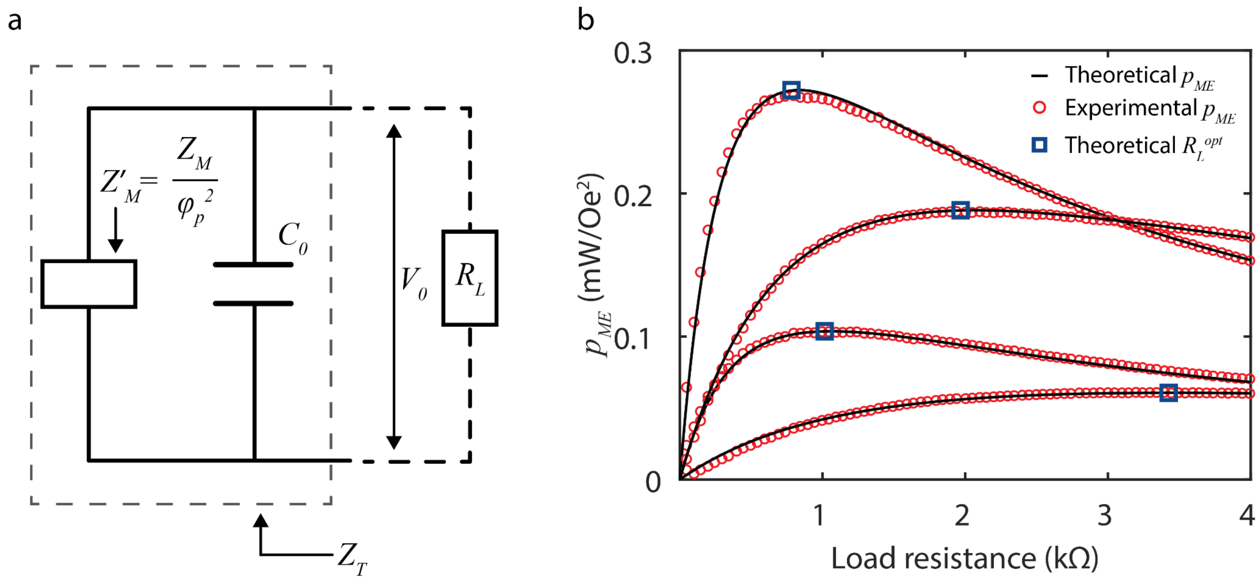


[FIG. S3. Calculating optimal load resistance from the equivalent circuit model. (a) The equivalent ME impedance viewed at the output terminal ($Z_T$) is expressed as the parallel combination of $Z'_M$ and $C_0$. Under the acoustic resonant frequency ($\omega_r$), the value of $R_L$ should match the real part of the $Z_T$ to receive maximum power. (b) Theoretical (black solid line) and experimental (red dots) value of $p_{ME}$ as a function of load resistance for four different version of ME receivers (Config. 1(top) - 4(bottom)) which analytical solution of $R_L^{opt}$ (blue squares) obtained from Eq. S30]

We can calculate the optimal load ($R_L^{opt}$) for the ME receiver under resonant frequency using the equivalent circuit model and predefined parameters. Based on the maximum transfer theorem, load impedance must be equal to the equivalent ME impedance as viewed from the output terminal ($Z_T$) to get maximum power from the source. As depicted in Fig. S3a, the equivalent ME impedance viewed at the output terminal ($Z_T$) can be defined as the parallel combination of the equivalent mechanical impedance reflected to the electric domain ($Z'_M = \frac{Z_M}{\varphi_p^2}$) and the capacitance of PZT ($C_0$) (Figure S3). Thus, $Z_T$ is expressed as:

$$Z_T = Z'_M || \frac{1}{j\omega C_0} = \frac{Z_M}{\varphi_p^2} || \frac{1}{j\omega C_0} = \frac{Z_M}{j\omega C_0 Z_M + \varphi_p^2}. \text{ (S27)}$$

At the acoustic resonant frequency, eq. S27 can be re-written by substituting $Z_M$ with the relation from Eq. S4 as:

$$Z_{T,\,\omega=\omega_r} = \frac{R_M + j\omega_r L_M + \frac{1}{j\omega_r C_M}}{j\omega_r C_0 (R_M + j\omega_r L_M + \frac{1}{j\omega_r C_M}) + \varphi_p^2}. \quad \text{(S28)}$$

Our previous study[7] has shown that the equivalent ME impedance at the resonant frequency is dominated by the resistive part of the impedance ($Im(Z_{T,\,\omega=\omega_r}) \simeq 0$). Therefore, we can regard the optimal load impedance as a resistor that matches with the real part of the equivalent ME impedance ($Re(Z_{T,\,\omega=\omega_r})$) as:

$$R_L^{opt} \simeq Re\left(Z_{T,\,\omega=\omega_r}\right) = Re\left(\frac{R_M + j\omega_r L_M + \frac{1}{j\omega_r C_M}}{j\omega_r C_0 (R_M + j\omega_r L_M + \frac{1}{j\omega_r C_M}) + \varphi_p^2}\right). \quad \text{(S29)}$$

By substituting variables in Eq. S29 with the expression from Eq. S2-S3 and S5-S12, we obtain $R_L^{opt}$ as:

$$R_L^{opt} \simeq Re\left(Z_{T,\,\omega=\omega_r}\right) = R(k, \eta) \times Q_M \times t_{total}, \quad \text{(S30)}$$

where $R(k, \eta) = \frac{8 d_{31,p}^2}{\varepsilon_r^2 \varepsilon_0^2 \pi^3 s_{11,p}^2} \eta^2 \left(\eta\rho_p + \frac{1-\eta}{k}\rho_m\right)^{1/2} \left(\frac{\eta}{s_{11,p}} + \frac{1-\eta}{k s_{33,m}}\right)^{-3/2}$.

Figure S3b shows the ME power coefficient from four different versions of ME receivers (config.1-4) as a function of load resistance. Red circles and solid lines represent the experimental and calculated values, respectively. Blue squares represent the analytical solution of optimal load resistance calculated with Eq. S30.

Figure S4 shows dependency of resistive part of the ME impedance, $Re(Z_{T,\,\omega=\omega_r})$, as a function of $Q_M$, $k$, $t_{total}$, and $\eta$. As depicted in Figs. S4, our model shows that $Re(Z_{T,\,\omega=\omega_r})$ is linearly related to the $Q_M$ and $t_{total}$. This results explain our finding that $p_{ME,\,max}$ is linearly related to $Q_M$ and $t_{total}$.

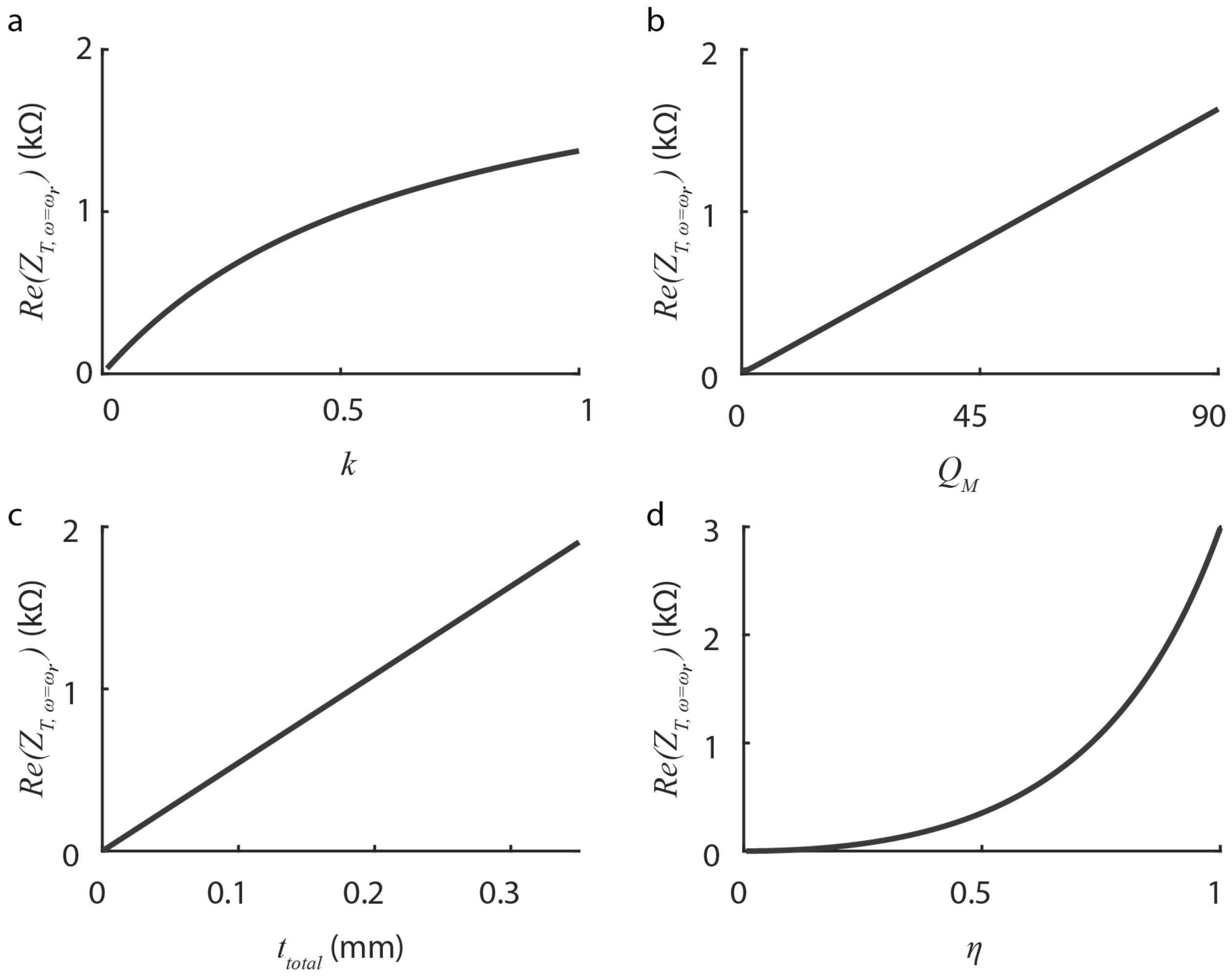


[FIG. S4. Plots show how the $Re(Z_{T,\,\omega=\omega_r})$ depends on several experimentally controlled variables based on Eq. S30 as a function of (a) $k$, (b) $Q_M$, (c) $t_{total}$, and (d) η. $Re(Z_{T,\,\omega=\omega_r})$ is linearly related to $Q_M$ and $t_{total}$ and non-linearly related to $k$ and η. ]

**Section 2. Parameter determination**

**2.1 Experimentally determining mechanical quality factor ($Q_m$)**

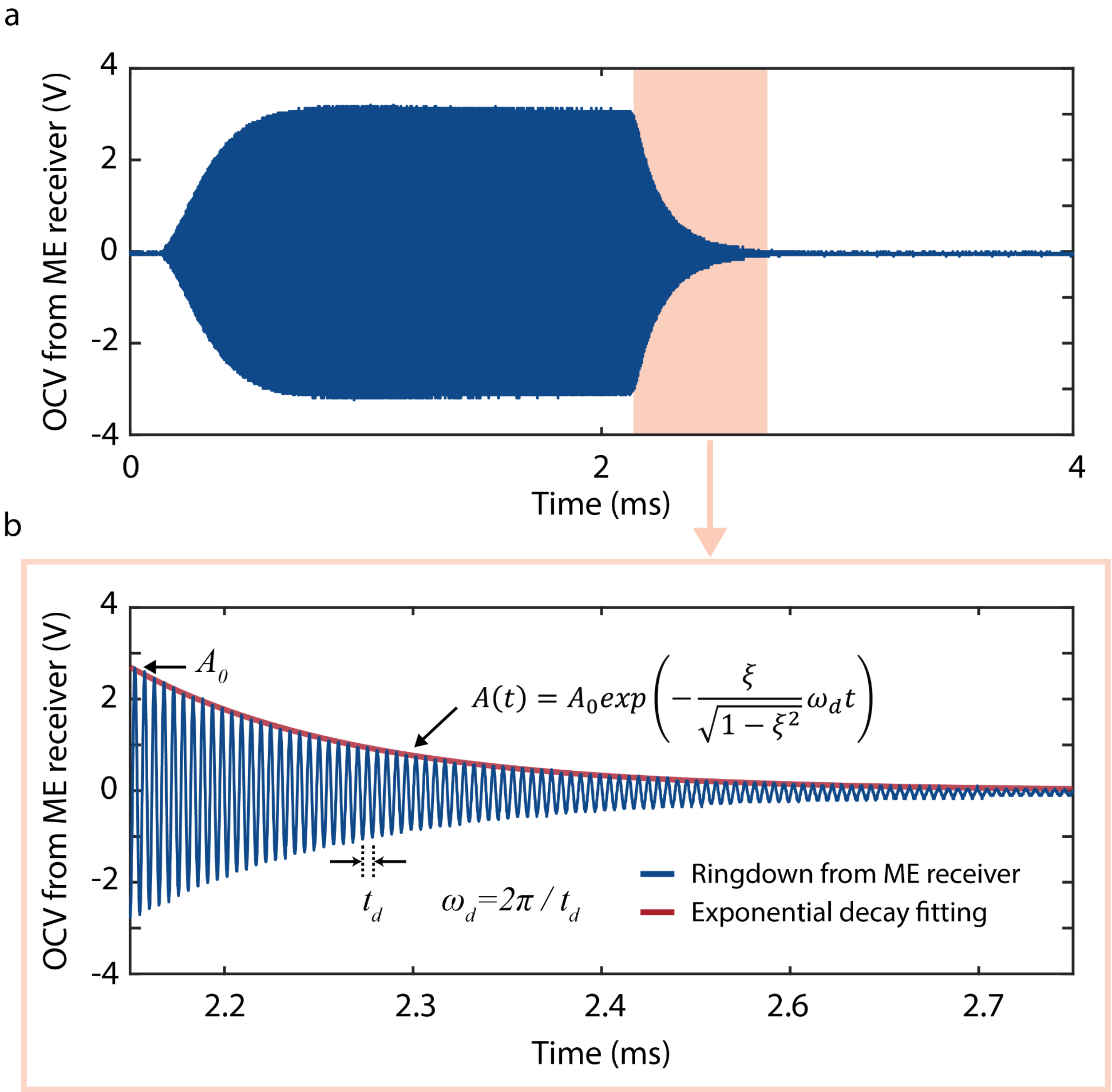


[FIG S5. Experimentally determining mechanical quality factor ($Q_M$) by fitting a ringdown signal from the ME receiver to the exponential decay envelope. (a) OCV from ME receiver at the acoustic resonant frequency. (b) Zoomed in version of Fig S4a to illustrate ring down signal from ME receivers (blue) and exponential decay fitting (red)]

The mechanical quality factor ($Q_M$) is defined as the ratio of the initial energy stored to the energy lost in one cycle[6]. The upper bound of $Q_M$ is constrained by the material property.

However this factor is also affected by the fabrication process and packaging. Thus, in this study, $Q_M$ of each ME receiver is determined experimentally. We can calculate the value of $Q_M$ assuming the ME transducer is a harmonic oscillator. The mechanical quality factor can be expressed as follows in terms of the damping ratio (ζ)[1,6]:

$$Q_M = 2\pi \frac{E_{stored}}{E_{lost\ per\ cycle}} = \frac{1}{2\zeta}. \quad (S31)$$

The OCV from the ME receivers shows exponential decay, generating the envelope shape (a ringdown) as depicted in Fig. S5. For an underdamped system ($0 < \zeta < 1$), the decaying amplitude of the ringdown can be expressed as:

$$A(t) = A_0 \exp\left(-\frac{\zeta}{\sqrt{1-\zeta^2}}\omega_d t\right), \quad (S32)$$

where $A_0$ and $\omega_d$ are initial amplitude and frequency of the ringdown, respectively. The damping ratio can be extracted with exponential fitting from the local maximum values of the ringdown as shown in Figure S5. The value of $Q_M$ is then determined by eq. S31. In this study, we were able to achieve $Q_M$ from 40 to 75.

**2.2 Experimentally determining interface coupling factor ($k$)**

The interface coupling factor ($k$) is defined as the ratio of the strain transferred from the magnetostrictive layer to the piezoelectric layer[1,7] ($0 < k < 1$). The value of interface coupling factor ($k$) depends on the material properties (i.e. Young's modulus) as well as the adhesion of the Metglas and PZT layers thus, should be experimentally determined. We can calculate this factor by fitting the experimental data to the theoretical model. The ME voltage coefficient as a function of $k$ is depicted in Fig.2a. The effect of $k$ changing $\omega_r$ follows relation in Eq. S5-S9 and S18 where $k$ affects average density and sound velocity, therefore, affecting characteristic impedance and frequency, finally affecting the resonant frequency $\omega_r$. Therefore, we can experimentally calculate $k$ by fitting the solution of $\omega_r$ from Eq. S18 to the experimental value. In this study, $k$ values ranged 0.42 ~ 0.70.

**Section 3. ME receiver fabrication and configurations**

For our experimental data collection, we used Metglas 2605SA1 from Metglas Inc. as the magnetostrictive layer and PZT-5A from Piezo Inc. as the piezoelectric layer to fabricate ME receivers. To ensure mechanical adhesion, we applied a thin layer of epoxy at the interface of the Metglas and PZT layers. The ME receivers were then laser cut to the desired shape and size using a custom-built femtosecond laser cutter. We fabricated ME receivers in two sizes, 5 × 2 $mm^2$ or 9 × 3 $mm^2$, and involved multiple researchers in the fabrication and testing process to reduce bias and human error.

Furthermore, we fabricated four different versions of ME receivers to validate the model validation and increase power by optimizing the total thickness and thickness ratio of the ME receivers. The configurations for these four versions are provided in Table SⅡ. We explicitly stated the size and configuration of the ME receivers that we used for each experiment to ensure transparency and reproducibility of our results.

[Table SⅡ. Configurations and thicknesses of four different ME receivers used in this study.]

| | Config. 1 | Config. 2 | Config.3 | Config.4 |
|---|---|---|---|---|
| 1st layer | Metglas | Metglas | Metglas | Metglas |
| 2nd layer | PZT | PZT | PZT | PZT |
| 3rd layer | Metglas | Metglas | - | - |
| $t_m$ (μm) | 50 | 50 | 25 | 25 |
| $t_p$ (μm) | 127 | 267 | 127 | 267 |
| $t_{total}$ (μm) | 177 | 317 | 152 | 292 |
| η | 0.72 | 0.84 | 0.84 | 0.91 |

**Section 4. Experimental setups**

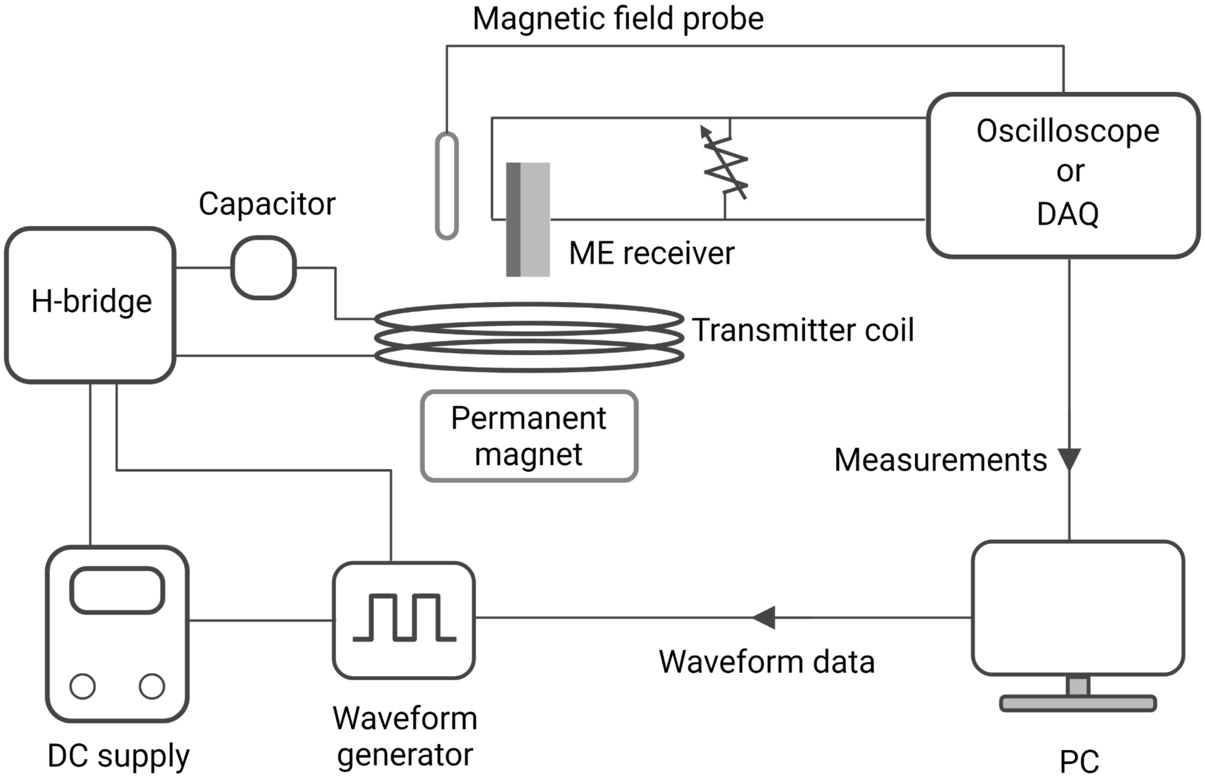


[Figure S6. Schematic of a benchtop setup to acquire waveform, voltage, and power measurements. We generated magnetic fields at desired strength and frequency using a computer, waveform generator, DC supply, H-bridge, capacitor, and transmitter coil. ME receivers were located at the center of the transmitter coil at a distance of 1-5 cm and a permanent magnet was manually placed at the optimum position according to the ME receiver's location. We used an oscilloscope or a DAQ for measurements. Acquired data was analyzed using MATLAB.]

We built a benchtop setup to acquire experimental data. We generated switching signals at desired frequency and duty cycle using a computer that is connected to the waveform generator (Analog Discovery2, Digilent). The signal is then transmitted to the H-bridge made up of four integrated driver ICs and MOSFETS (CSD95378BQ5M, Texas Instruments). The H-bridge features 30 Amps continuous current with a frequency limit of 1.25MHz. The H-bridge is connected to the transmitter coil which is wrapped with 40 AWG Litz Wire (MWS Wire) having an inductance of 25.6 μH. The transmitter coil is serially connected to the high-voltage rated film capacitors (WIMA GmbH & Co) to form a resonant circuit at desired driving frequency. The ME receiver is placed 1-5 cm above the surface of the transmitter coil and aligned at the center of the coil such that its longitudinal axis is perpendicular to the surface of the transmitter coil. A permanent magnet is used to provide a DC bias field to enhance the ME performance and was manually placed at the optimum position where the ME receiver showed its maximum OCV.

We acquired OCV and waveform data from the ME receivers using an oscilloscope (DS1104, Rigol). Power measurements from the ME receiver were obtained using a custom resistance sweeping system that we designed and connected to a DAQ (cDAQ-9171, NI). This system is controlled by a Teensy LC microcontroller, enabling us to read the voltage across the resistance while sweeping the load resistance from 50Ω to 4kΩ, with a resolution of 50Ω. We analyzed the acquired data using MATLAB. To measure the strength of the magnetic field, we utilized a 2D HF Magnetic Field Probe (AMF Lifesystems). The B-field was measured using a magnetic field probe, and we subsequently converted it to the corresponding H-field measurement to match the input component of the equivalent circuit model. To perform the conversion, we employed the relation $H = B/\mu_0$, where $\mu_0$ denotes vacuum permeability. We utilized the relation $1\ A/m = 4\pi \times 10^{-3}\ Oe$ for the unit conversion in H-field.

**Section 5. High power delivery demonstration through porcine tissue**

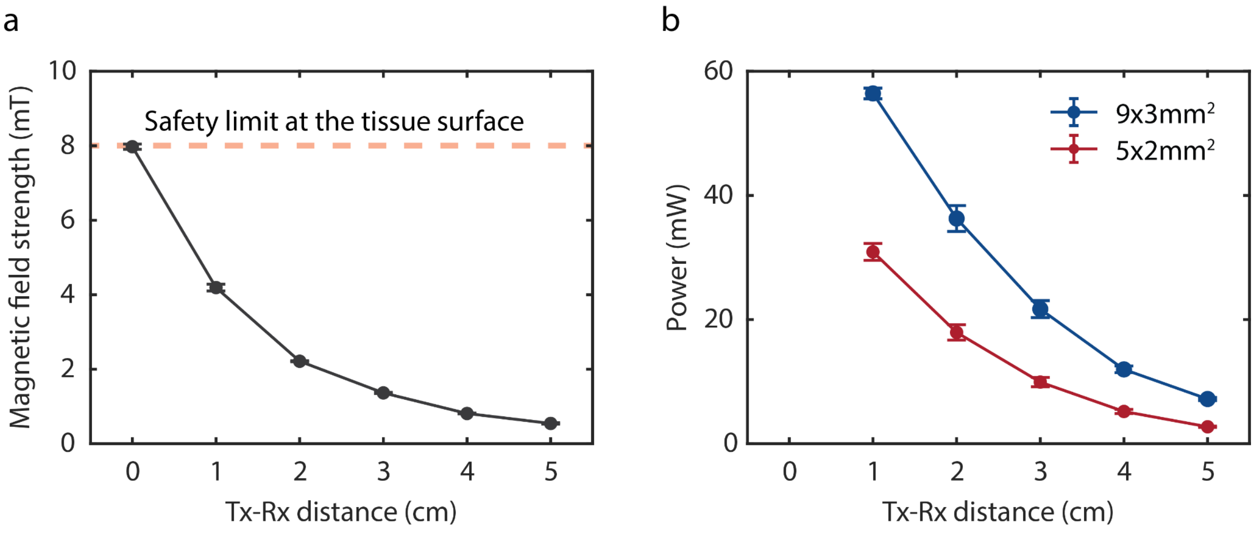


[Figure S7. Results of high power delivery demonstration through *ex vivo* porcine tissue. (a) We measured magnetic field strength at the surface of the tissue (0 cm) and through porcine tissue (1 to 5 cm). The orange line expresses the safe magnetic field strength limit for humans at the skin layer in unrestricted environments (8 mT). We ensured that the magnetic field strength at the surface of the porcine tissue remained below this safe limit of 8 mT. Measured magnetic field strength decreased from 4.2 to 0.5 mT as the distance increased from 1 to 5 cm, respectively. (b) We measured power in two different sized ME receivers as a function of distance through tissue (n=3 for each size). Received powers from 9 × 3 $mm^2$ ME receivers (blue) and 5 × 2 $mm^2$ ME receivers (red) ranged from 56.4 to 7.2 mW and 31.0 to 2.7 mW, respectively, at distances ranging from 1 to 5 cm.]

We have demonstrated that our optimized ME receivers can deliver high power density in *ex vivo* porcine tissue while adhering to human safety limits. To achieve this, we adjusted the input current of the transmitter coil to maintain a magnetic field strength of 8.0 mT at the surface of the porcine tissue. This value is within the permissible human safety limits in unrestricted environments determined by our previous study[7]. To further investigate the capabilities of our system, we placed 1cm thick slices of porcine tissue between the transmitter coil and ME receivers to simulate implantation depths ranging from 1 to 5 cm. We then measured both the magnetic field strength and received power at each distance. Our results showed that the measured magnetic field strength decreased from 4.2 to 0.5 mT as the distance increased from 1 to 5 cm, respectively (Fig. S7a). Additionally, we measured the power output of two different sized ME receivers: 9 × 3 $mm^2$ and 5 × 2 $mm^2$. With the 9 × 3 $mm^2$ ME receivers, we achieved a power output range of 56.4 to 7.2 mW at distances ranging from 1 to 5 cm, respectively (Fig. S7b, blue). Similarly, we achieved a power output range of 31.0 to 2.7 mW with the 5 × 2 $mm^2$ ME receivers at the same distances (Fig. S7b, red).

**Section 6. Comparison of this work with other WPT modalities**

We compared power delivery of our optimized ME receivers with similarly sized bioelectronic devices powered by ME, near-field inductive coupling (NIC), light, radiofrequency electromagnetic waves (RF), ultrasound (US). More details can be found in Table SⅢ.

[Table SⅢ. Comparison with other WPT modalities in similar size and implantation depth.]

| **Reference** | this work | this work | A[8] | B[9] | C[10] | D[11] | E[12] | F[13] | G[14] | H[15] | I[16] |
|---|---|---|---|---|---|---|---|---|---|---|---|
| **Modality** | ME | ME | ME | NIC | NIC | NIC | light | US | US | RF | MDF |
| **Area of receiver ($mm^2$)** | 10.00 | 27.00 | 10.00 | 0.09 | 16.00 | 32.00 | 0.24 | 0.36 | 19.50 | 2.56 | 4.50 |
| **Power (mW)** | 30.72 | 56.48 | 2.00 | 1.00 | 10.00 | 5.00 | 2.30 | 3.00 | 3.00 | 1.21 | 1.00 |
| **Power density ($mW/mm^2$)** | 3.07 | 2.09 | 0.20 | 11.11 | 0.63 | 0.16 | 9.58 | 8.33 | 0.15 | 0.47 | 0.22 |
| **Length (mm)** | 5.00 | 9.00 | 5.00 | 0.30 | 4.00 | 16.00 | 0.4 (single unit) | 0.60 | 3.00 | 1.6 | 3.00 |
| **Width (mm)** | 2.00 | 3.00 | 2.00 | 0.30 | 4.00 | 2.00 | 0.4 (single unit) | 0.60 | 6.50 | 1.6 | 1.50 |
| **Tx-Rx distance (cm)** | 1.0 | 1.0 | 2.5 | 0.4 | 1.1 | NA | NA | 10.0 | 10.5 | 1.0 | 4.6 |
| **Demonstration medium** | porcine tissue | porcine tissue | mice | air | ovine tissue | NA | air | mineral oil | castor oil | galline tissue | porcine tissue |
